# Integrating Image and Molecular Profiles for Spatial Transcriptomics Analysis

**DOI:** 10.1101/2023.06.18.545488

**Authors:** Xi Jiang, Shidan Wang, Lei Guo, Zhuoyu Wen, Liwei Jia, Lin Xu, Guanghua Xiao, Qiwei Li

## Abstract

The spatially resolved transcriptomics (SRT) field has revolutionized our ability to comprehensively leverage image and molecular profiles to elucidate spatial organization of cellular microenvironments. Current clustering analysis of SRT data primarily relies on molecular information and fails to fully exploit the morphological features present in histology images, leading to compromised accuracy and interpretability. To overcome these limitations, we have developed a multi-stage statistical method called iIMPACT. It includes a finite mixture model to identify and define histology-based spatial domains based on AI-reconstructed histology images and spatial context of gene expression measurements, and a negative binomial regression model to detect domain-specific spatially variable genes. Through multiple case studies, we demonstrate iIMPACT outperformed existing methods, confirmed by ground truth biological knowledge. These findings underscore the accuracy and interpretability of iIMPACT as a new clustering approach, providing valuable insights into the cellular spatial organization and landscape of functional genes within spatial transcriptomics data.

---

Spatially resolved transcriptomics (SRT), a new generation of RNA-sequencing analysis techniques, provides biological information at the cellular level while preserving the organization of the tissue and cellular microenvironment ^1, 2, 3, 4^. One category of SRT methods builds upon next-generation sequencing (NGS)-based SRT techniques, including spatial transcriptomics (ST) ^5^, 10x Visium (an improved ST platform), Slide-seq ^6^, Slide-seqV2 ^7^, and high-definition spatial transcriptomics (HDST) ^8^. These techniques capture RNA molecules via spatially arrayed barcoded probes. The barcoded areas, namely spots, cover a group of cells and are usually arrayed on a two-dimensional grid. Another category of SRT platforms is based on imaging techniques, such as seqFISH ^9^, MERFISH ^10^, and STARmap ^11^. They measure the expression level for hundreds to thousands of genes at the single-cell resolution with detailed spatial organization information. With these advancements, SRT techniques have been widely applied to facilitate discoveries of novel insights in biomedical studies.

A central challenge for SRT data analysis is to define clinically or biologically meaningful spatial domains by partitioning regions with similar molecular and/or histological characteristics, because the spatial domain identification serves as the foundation for several important downstream analyses, including but not limited to the domain-based differential expression analysis, trajectory analysis, and functional pathway analysis ^12, 13^. However, current state-of-the-art methods typically focus on achieving this goal solely by analyzing SRT molecular profiles, such as gene expressions, while neglecting the valuable morphological or biological information present in the associated histology images. For example, the Seurat package, the most prevalent single-cell RNA sequencing data analysis pipeline ^14, 15^, utilizes only the high-throughput gene expression of each spot for clustering analysis but does not leverage any information from the associated histology images in this analysis. On the other hand, several recently developed methods, such as stLearn ^16^, hidden-Markov random field ^17^, BayesSpace ^18^, and SpaGCN ^19^, integrate spatial information or various features extracted from the histology image into the clustering analysis of SRT data. However, those image features, such as RGB color values, do not explicitly reveal detailed morphological information (e.g., cell locations and types) and therefore fail to provide biologically relevant insights.

Different from molecular information, histology images characterize cellular structures and tissue microenvironments, which have been proven valuable in clinical diagnosis and prognosis^20, 21^. Computer vision algorithms have enabled us to automatically segment cell nuclei from digital histology images at a large scale ^22^. Recent developments in deep convolutional neural networks (e.g., H-DenseUNet ^23^, Micro-Net ^24^, Hover-Net ^25^, and HD-Staining model ^20^) have further integrated the automatic identification, classification, and feature extraction of each observed nucleus in a histology image. In practice, a histology-based spatial domain (e.g., tissue) is defined as a group of cells with similar morphological and molecular context as a unit. Thus, we hypothesize that integrating spot-level molecular profiles and cellular-level image profiles from AI-reconstructed histology images could enhance the spatial domain identification in terms of both accuracy and interpretability.

Another challenge for SRT data analysis is to identify spatially variable genes (SVGs), which represent genes with spatially correlated expression patterns ^26, 27^. Recently developed methods, such as SpatialDE ^28^, SPARK ^26^, BOOST-GP ^27^, and BOOST-MI ^29^, characterize the global spatial dependency of a gene in the whole domain while ignoring the spatial pattern heterogeneity due to cellular organization, which could be fully observed in AI-reconstructed histology images. SpaGCN ^19^ proposed domain-guided differential expression analysis to detect SVGs without a rigorous statistical framework. Therefore, there is an urgent need to develop reliable statistical method to detect domain-specific SVGs.

This paper proposes a two-stage statistical approach by integrating Image and Molecular Profiles to Analyze and Cluster spatial Transcriptomics data, or iIMPACT for short. The first stage is implementing a Bayesian finite mixture model to allocate all spots into mutually exclusive clusters, namely histology-based spatial domains. We decompose each mixture component into two sub-components to integrate image and molecular profiles. In particular, a multinomial sub-component is employed to model cell type abundance available in histology images. Following BayesSpace ^18^, we use a normal sub-component to model the low-dimensional representation of normalized gene expression from the matching SRT molecular profile. The Bayesian model also adopts a Markov random field prior (MRF) to encourage neighboring spots to be clustered in the same histology-based spatial domain. The spots’ neighborhood structure can be straightforwardly defined from the NGS-based SRT geospatial profile, as spots are usually arrayed on square or triangular lattices. Through the resulting posterior inference, we obtain histology-based spatial domains and their interactive zones, while characterizing each identified histology-based spatial domains by inferring its underlying domain-specific relative abundance of cell types. The second stage is implementing a negative binomial (NB) regression model to search for domain-specific SVGs, which are differentially expressed between a given histology-based spatial domain identified in the first stage and all others. This approach directly models the numbers of read counts (used as a proxy for gene expression) in the SRT molecular profile to achieve minimum information loss. iIMPACT could also be extended to analyze imaging-based SRT data via some special handling. Compared with existing state-of-the-art methods, iIMPACT is able to fully leverage information from the nuclei segmentation procedure on the histology images for clustering analysis and has strong biological interpretability. Applying iIMPACT on four datasets from different SRT platforms (summarized in Table S1), we confirmed that iIMPACT performed better on both spatial domain identification and domain-specific SVG detection than state-of-the-art methods. We further demonstrated that iIMPACT could capture biological features at both the spatial domain level and gene level. Therefore, by integrating image and molecular information, iIMPACT facilitates the discovery of new biological insights from SRT datasets.

## RESULTS

### Overview of iIMPACT

iIMPACT is a two-stage statistical method to analyze SRT data, with its workflow shown in Figure 1. It includes two stages – histology-based spatial domain identification by a Bayesian normal-multinomial mixture model and domain-specific SVG detection by an NB regression model.

**Figure 1.**
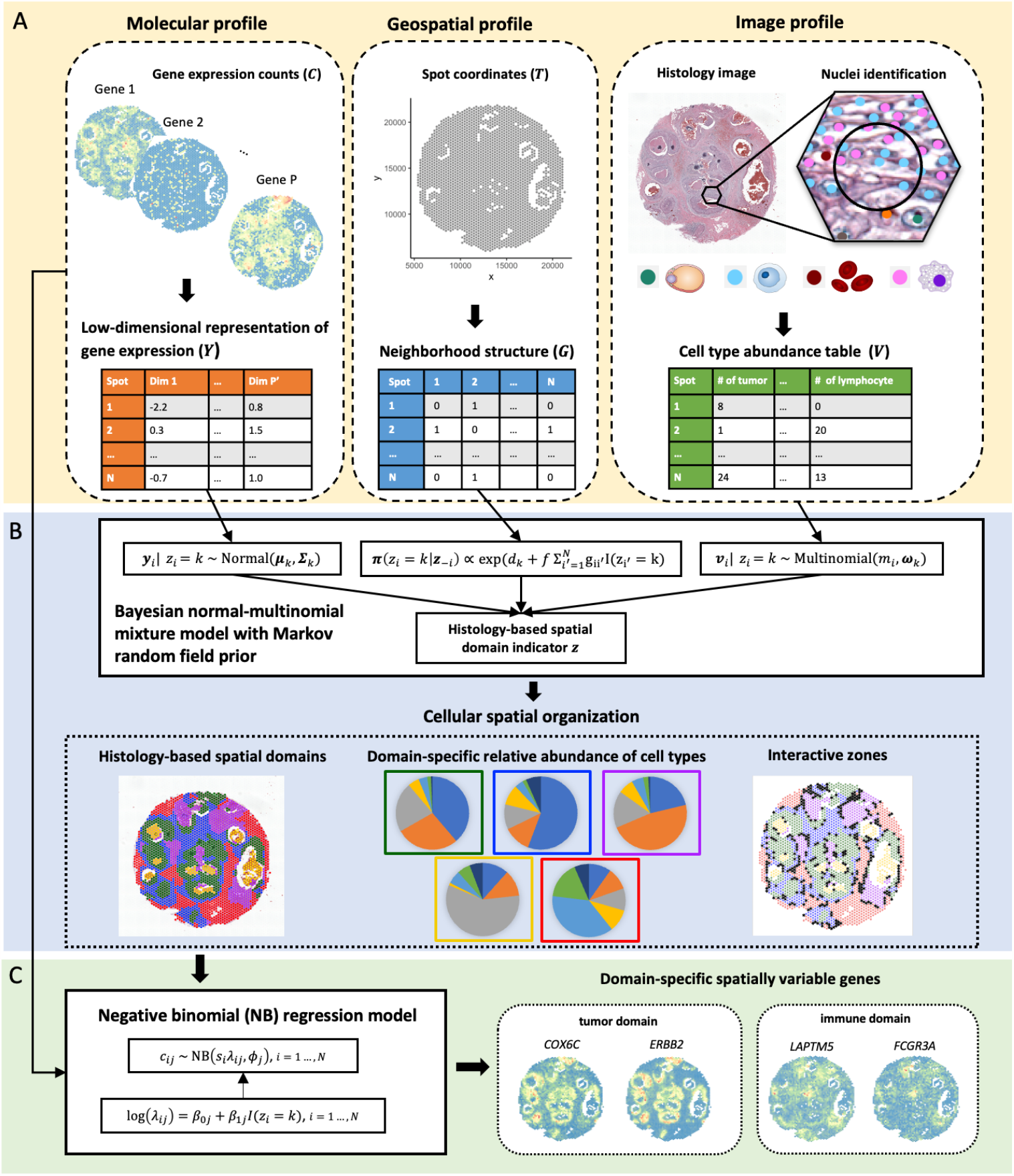
Workflow of iIMPACT: A. iIMPACT starts by combining and processing image profile from AI-reconstructed histology images, and geospatial and molecular profiles from SRT data (circled by dashed lines) to conduct the histology-based spatial domain identification. B. A Bayesian normal-multinomial mixture model with the Markov random field (circled by solid lines) is fitted for histology-based spatial domain identification. Based on the spatial domain identification results, biologically important cellular spatial organization can be characterized, including the domain-specific relative abundance of cell types and interactive zones (circled by dotted lines). C. Domain-specific SVGs are identified by a negative binomial (NB) regression model.

To achieve the above goals, iIMPACT utilizes the morphological context of histology images and the spatial context of gene expression measurements, referring to the image and molecular profiles in Figure 1A and throughout the paper. In particular, the molecular profile refers to the low-dimensional representation of normalized gene expression values at the spot level (denoted by ***Y***), which is obtained by a pre-specified dimension reduction technique, such as principal component analysis (PCA). The accompanying SRT geospatial profile that records all spots’ locations is processed as an adjacent matrix (denoted by ***G***) representing the spots’ neighborhood structure. iIMPACT requires the locations and types of all cell nuclei in the matching histology image. Combining with the geospatial profile, we can generate the image profile (denoted by ***V***), which indicates the spot-level cell type abundance, i.e., the numbers of different cell types within a spot and its expanded area.

In the first stage, we employ a Bayesian normal-multinomial mixture model with the MRF prior ^30, 31^ to identify the histology-based spatial domains (Figure 1B) and interactive zones, corresponding to those spots with less confidence to be allocated to any histology-based spatial domains. Through model parameter estimation, iIMPACT can infer the underlying relative abundance of cell types at each histology-based spatial domain to provide a reference to distinguish their histological types. In the second stage, an NB regression model is fitted for each gene and each histology-based spatial domain of interest, where domain-specific SVGs can be defined (Figure 1C).

### Application to human breast cancer dataset

We applied iIMPACT to analyze an SRT dataset from a human breast cancer study. This dataset includes 2,518 spots and 17,651 genes. The gene expression was measured on a section of human breast with invasive ductal carcinoma via the 10x Visium platform, along with annotation from pathologists that was used as the ground truth for algorithm comparison purposes (H&E-stained image with five annotated tissue regions in Figure 2A). After applying HD-Staining ^20^ to the histology image of breast cancer tissue, we identified 156,235 cells within seven categories: macrophage, ductal epithelium, karyorrhexis, tumor cell, lymphocyte, red blood cell, and stromal cell (Detailed information in Figure S1).

**Figure 2.**
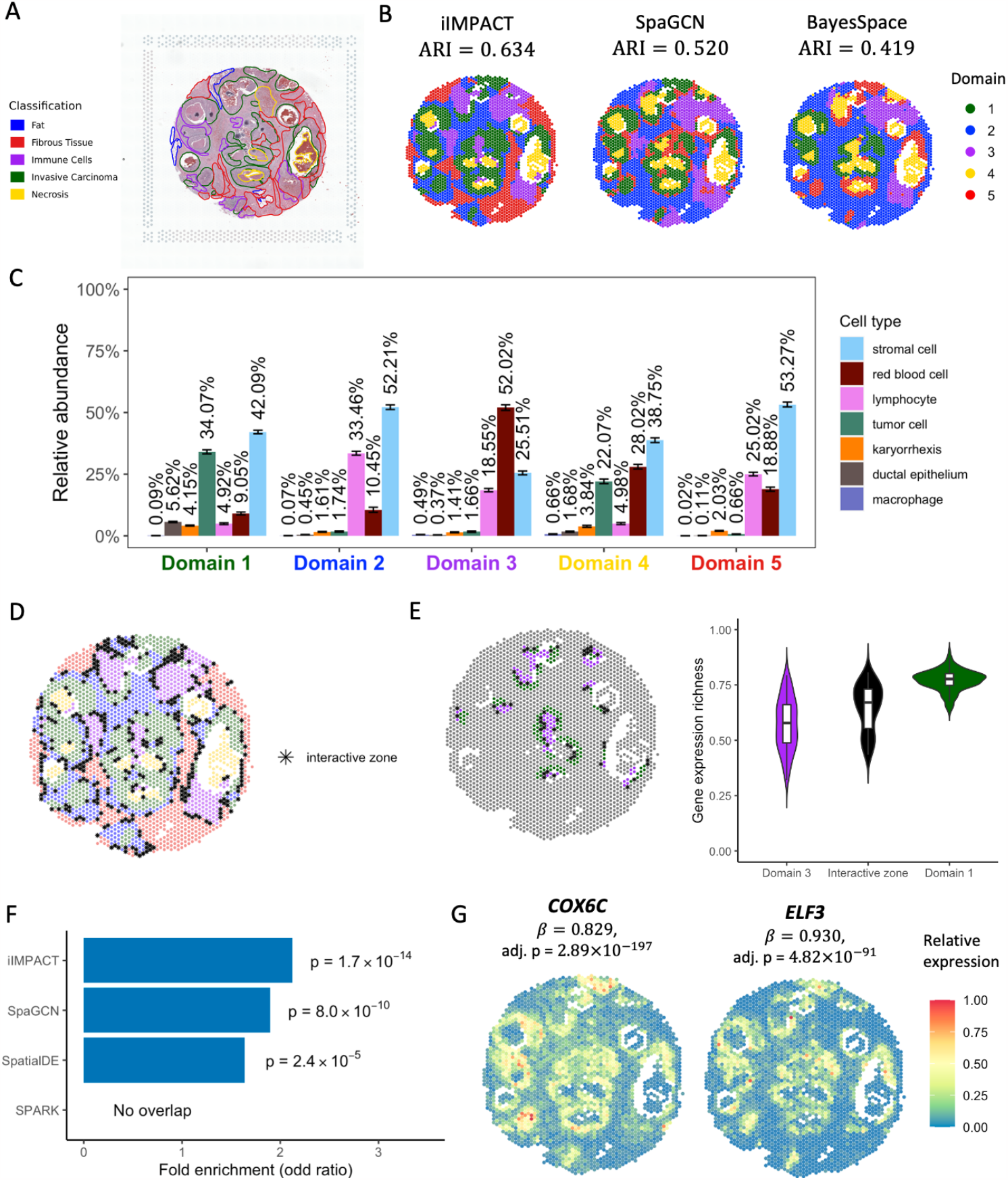
Human breast cancer dataset: A. H&E-stained image of the tissue section with ground truth labels from pathologists. B. Spatial domains detected by iIMPACT, SpaGCN, and BayesSpace, with the number of clusters to be five. C. Estimates (posterior means and credible intervals) of domain-specific relative abundance of cell types for the seven cell types observed in the AI-reconstructed histology image. D. Interactive zones (black asterisk spots) defined by iIMPACT. E. Identified interactive zones (black asterisk spots) and other boundary areas of tumor and its adjacent Domain 3, and boxplots of gene expression richness for spots in the interactive zone and other boundaries. F. Gene enrichment analysis between SVGs detected by iIMPACT, SpaGCN, SpatialDE, and SPARK, and known breast cancer genes from the COSMIC database. G. Spatial expression patterns of two example SVGs, *COX6C* and *ELF3*, that were only detected by iIMPACT.

Firstly, we compared the five spatial domains defined by iIMPACT, SpaGCN ^19^, and BayesSpace ^18^, with manually annotated domains by pathologists. We found that iIMPACT achieved the highest consistency with the manual annotation (See Figure 2B. Adjusted Rand Index (ARI) = 0.634), while BayesSpace mistakenly clustered the tumor region into two groups. It is worth noting that SpaGCN took only the image RGB values instead of detailed histology information, which might contribute to the unsatisfactory performance on segmenting the non-tumor regions (ARI = 0.520). In contrast, the better performance of iIMPACT suggests the advantage of integrating both molecular and image profiles in the clustering analysis of SRT data.

Secondly, iIMPACT is able to define each individual histology-based spatial domain through its underlying relative abundance of cell types parametrized by the Bayesian multinomial-normal mixture model (Figure 2C). In contrast, SpaGCN and BayesSpace, despite their good capabilities in identifying spatial domains, currently lack the ability to effectively integrate cell type information and interpret the identified domains in a biologically meaningful way. For example, as detailed in Figure 2C, the proportion of tumor cells is higher in domain 1 (green spots in Figure 2B) than in other domains, indicating that domain 1 is the tumor region. This inference is consistent with tumor regions in the manual annotation. Domain 2 (blue) and domain 5 (red) have a similar proportion of stromal cells, while the proportion of lymphocytes in domain 2 is higher than in domain 5. The difference in the relative abundance of cell types may indicate the functional difference between these two domains. These examples confirm that iIMPACT is able to provide biological interpretation of spatial domains.

Thirdly, iIMPACT can identify the interactive zones among histology-based spatial domains (Figure 2D). Interactive zones are spots with higher uncertainty on domain allocation, which potentially have higher diversity in cell type abundance and heterogeneity in gene expression compared with neighboring spots with unambiguous domain definition. We calculated the gene expression richness, defined as the percentage of genes with non-zero read counts, for each spot in the tumor-immune interactive zone and other tumor-immune boundaries. We observed statistically significant differences among these comparisons (Figure 2E), implying that the identified zones are connected areas between tumor and immune domains with a high level of heterogeneity in gene expression and complex cellular interactions. By further comparing the gene expressions for these groups, we found several known cancer or immune genes with high expression in the interactive zones (e.g., *GREM1* ^*32*^), suggesting the possible tumor-immune interactions in these zones.

Finally, we asked whether the domain-specific SVGs defined by iIMPACT are more consistent with biological knowledge than those from other algorithms, which is an independent evaluation step frequently used for validating the clustering approaches on single-cell and spatial profiling data ^19, 26, 28^. We focused on the tumor-domain specific SVGs defined by iIMPACT, SpaGCN ^19^, SpatialDE ^28^, and SPARK ^26^, respectively, and performed the enrichment analysis by comparing tumor-domain SVGs defined by these four methods with the known breast cancer gene set defined in the Catalogue Of Somatic Mutations In Cancer (COSMIC) database. As summarized in Figure 2F, the tumor-domain SVGs detected by iIMPACT showed higher overlap with the known breast cancer gene set than that of SpaGCN, SpatialDE, and SPARK, respectively, including two example genes that can only be detected by iIMPACT (Figure 2G): *COX6C* (estimated coefficient 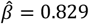, with adjusted p-value = 2.89 × 10^−197^), a known biomarker for the identification of hormone-responsive breast cancer ^33^, and *ELF3* (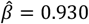, with adjusted p-value = 4.82 × ^−91^), an epithelial-specific gene that is a novel therapeutic target of breast cancer and has been amplified in early breast cancer ^34^. These results confirm that iIMPACT-defined SVGs align better with existing biological knowledge.

### Application to human prostate cancer dataset

To evaluate the performance of iIMPACT in different tissue types, we studied another SRT dataset from a human prostate cancer study, which includes 4,371 spots and 17,651 genes. The gene expression was measured on a section from invasive carcinoma of human prostate via the 10x Visium platform. We applied HD-Staining to analyze the histology image of this tissue (Figure 3A). 259,257 cells were segmented and classified into six categories: macrophage, karyorrhexis, tumor cell, lymphocyte, red blood cell, and stromal cell (Detailed information in Figure S2).

**Figure 3.**
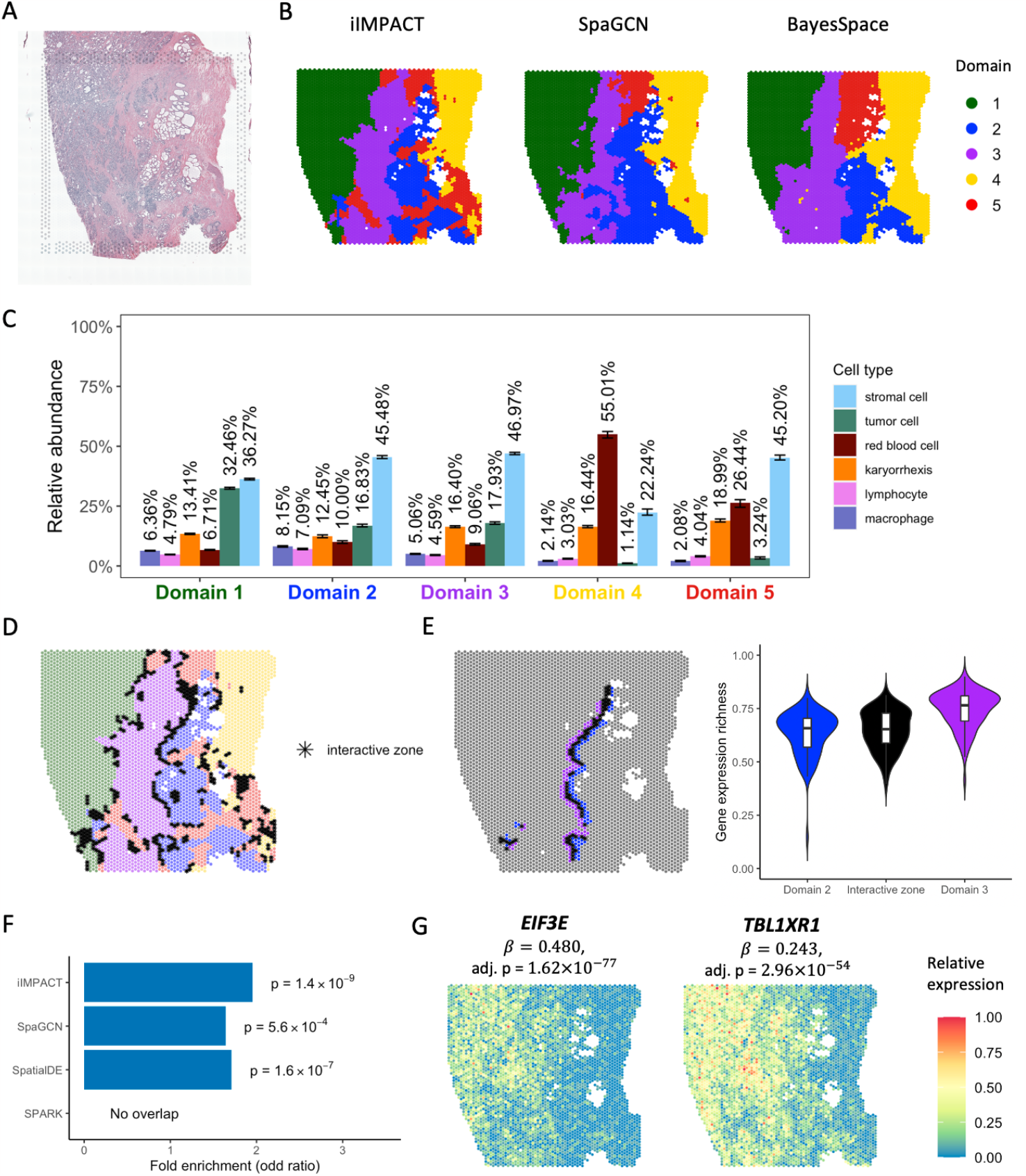
Human prostate cancer dataset: A. H&E-stained image of the tissue section. B. Spatial domains detected by iIMPACT, SpaGCN, and BayesSpace, setting the number of clusters to be five. C. Estimates (posterior means and credible intervals) of domain-specific relative abundance of cell types for the six cell types observed in the AI-reconstructed histology image. D. Interactive zones (black asterisk spots) defined by iIMPACT. E. Identified interactive zones (black asterisk spots) and other boundary areas of Domain 2 and Domain 3, and boxplots of gene expression richness for spots in the interactive zone and other boundaries. F. Gene enrichment analysis between SVGs detected by iIMPACT, SpaGCN, SpatialDE, and SPARK, and the known prostate cancer genes from the COSMIC database. G. Spatial expression patterns of two example SVGs, *EIF3E* and *TBL1XR1*, that were only detected by iIMPACT.

We confirmed that iIMPACT outperformed BayesSpace and SpaGCN in spatial domain detection, assuming there are five spatial domains. As shown in Figure 3B, these three methods could identify the domain (marked in green) with a high proportion of tumor cells, compared with the spatial distribution of tumor cells (Figure S2C). Interestingly, iIMPACT could distinguish histology-based spatial domains with different red blood cell proportions (Figure 3B, yellow region vs. red region). We further compared the results of three methods with the manual annotation based on three morphologically distinguish regions: tumor, stroma and partially atrophic changes, and stroma (Figure S3). We observed that iIMPACT achieved the highest consistency with the manual annotation (ARI = 0.659).

To demonstrate the interpretability of iIMPACT, we characterized the domain-specific relative abundance of cell types in Figure 3C. We observed that domain 1 has a higher proportion of tumor cells than other domains, indicating that it is probably the tumor domain. Comparing domain 2 with domain 3, we observed that they have a similar proportion of tumor cells, but domain 2 has a higher proportion of immune cells (i.e., lymphocyte and macrophage), implying the heterogeneity of immune composition within tumors.

In addition, interactive zones can also be defined by iIMPACT (Figure 3D). By checking the interactive zones of domain 2 and 3 and calculating the gene expression richness, we observed a clear trend between the interactive zones and the surrounding boundaries, indicating the unique characteristics of interactive zones (Figure 3E). We further found that gene *DNAJC5* ^35^ expressed higher on the identified interactive zones, implying its potential relationship with the intermediate areas of immune cell distribution.

We also compared iIMPACT, SpaGCN ^19^, SpatialDE ^28^, and SPARK ^26^ in detecting biologically meaningful SVGs in this prostate cancer dataset. We confirmed that, for tumor-domain (domain 1) specific SVGs, iIMPACT outperformed SpaGCN, SpatialDE, and SPARK in detecting known prostate cancer genes from the COSMIC database (Figure 3F), illustrating that iIMPACT could detect SVGs that are biologically relevant. These iIMPACT-defined SVGs in tumor domains have experimental evidence to support their functional relevance to the development of prostate cancer.

For example, as shown in Figure 3G, *EIF3E* (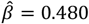, with adjusted p-value = 1.62 × 10^−77^), which is associated with increased cell cycle progression and motility in prostate cancer ^36^, and *TBL1XR1* (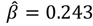, with adjusted p-value = 2.96 × 10^−54^), which displays an oncogene role for prostate cancer cell proliferation ^37^.

### Application to human ovarian cancer dataset

The third NGS-based SRT dataset is from a section of human ovarian tumor tissue. This dataset includes 3,455 spots and 17,651 genes. The gene expression was measured on a section of serous papillary carcinoma from human ovarian via the 10x Visium platform, with the H&E-stained image shown in Figure 4A, HD-Staining model segmented and classified 211,746 cells in six categories: macrophage, karyorrhexis, tumor cell, lymphocyte, red blood cell, and stromal cell (Detailed information in Figure S4). By utilizing the cell type abundance information from the histology image, we observed that iIMPACT had better performance on spatial domain identification. By comparing the clustering results of three methods (iIMPACT, BayesSpace, and SpaGCN) with the annotated tumor and benign domains for this SRT dataset, we observed a remarkable concordance between the clustering results obtained from iIMPACT and the pathologist’s annotations (ARI = 0.967, see Figure S5). In addition, iIMPACT could identify the domain (marked in green) with a high proportion of tumor cells, which has a high consistency with the tumor region annotated by the pathologist (Figure S5) and the region with a high amount of tumor cells (Figure S4C), while the other two methods could not confirm it as a tumor domain (Figure 4B).

**Figure 4.**
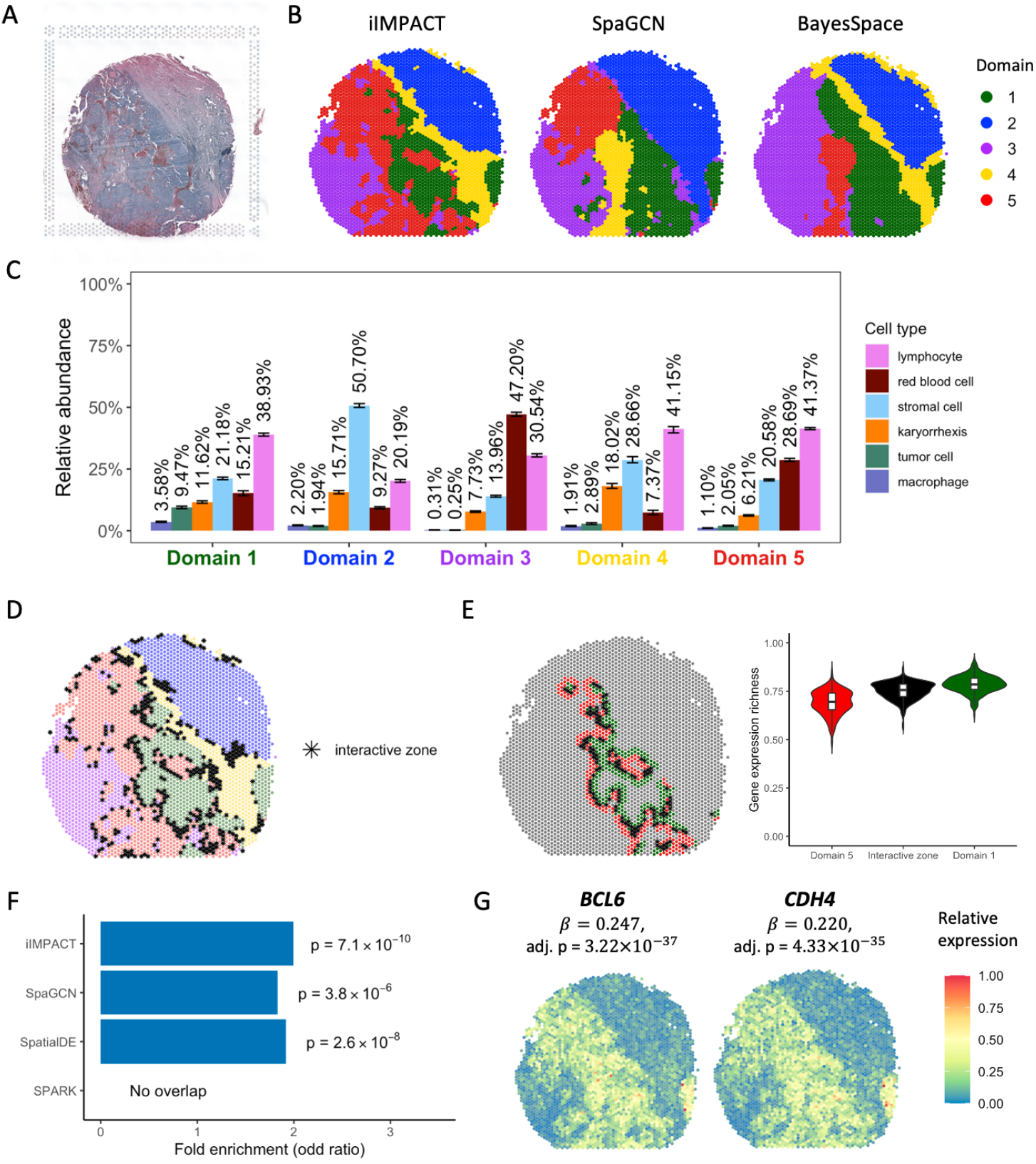
Human ovarian cancer dataset: A. H&E-stained image of the tissue section. B. Spatial domains detected by iIMPACT, SpaGCN, and BayesSpace, setting the number of clusters to be five. C. Estimates (posterior means and credible intervals) of domain-specific relative abundance of cell types for the six cell types observed in the AI-reconstructed histology image. D. Interactive zones (black asterisk spots) defined by iIMPACT. E. Identified interactive zones (black asterisk spots) and other boundary areas of tumor and its adjacent Domain 5, and boxplots of gene expression richness for spots in interactive zone and other boundaries. F. Gene enrichment analysis between SVGs detected by iIMPACT, SpaGCN, SpatialDE, and SPARK, and the known ovarian cancer genes from the COSMIC database. G. Spatial expression patterns of two example SVGs, *BCL6* and *CHD4*, that were only detected by iIMPACT.

iIMPACT could also distinguish domains with different red blood cell proportions. Figure 4C shows the estimation of relative abundance of cell types for the five histology-based spatial domains. Domain 1 has a higher proportion of tumor cells than other domains, indicating that it is likely to be the tumor domain. We further examined the interactive zones (Figure 4D) and compared the interactive zone between domain 1 and 5 with other boundary spots (Figure 4E). A significant difference in gene expression richness between boundary spots and the interactive zone was observed. Furthermore, we found that gene *TTLL5* ^38^ and *CLEC12A* ^39^ have higher expression on the interactive zone between domain 1 and 5, which may infer their potential relationship with the tumor-immune interaction.

We further detected SVGs using iIMPACT, and then queried tumor-region SVGs with the known ovarian cancer gene set defined by the COSMIC database. We observed that iIMPACT-defined ovarian cancer SVGs showed a higher overlap with the known ovarian cancer gene set than that that of SpaGCN, SpatialDE, and SPARK (Figure 4F). Moreover, we explored these ovarian cancer SVGs only defined by iIMPACT and found that many of them possess compelling experimental evidence substantiating their functional relevance to ovarian cancer. For example, our list included *BCL6* (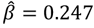, with adjusted p-value = 3.22 × 10^−37^), which displays pro-oncogenic activity in ovarian cancer ^40^, and *CHD4* (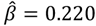, with adjusted p-value = 4.33 × 10^−35^), which is associated with apoptosis mediated by cisplatin in ovarian cancer cells ^41^ (Figure 4G).

### Application to mouse visual cortex STARmap data

To demonstrate iIMPACT is also able to analyze data from imaging-based SRT platforms, we applied iIMPACT to a STARmap dataset ^11^. This dataset was generated from mouse visual cortex, including hippocampus, corpus callosum, and the neocortical layers. In total, 1,020 genes were measured in 1,207 cells with 15 cell types. The layer structure and cell type distribution of the tissue section provided by the original study are displayed in Figure 5A.

**Figure 5.**
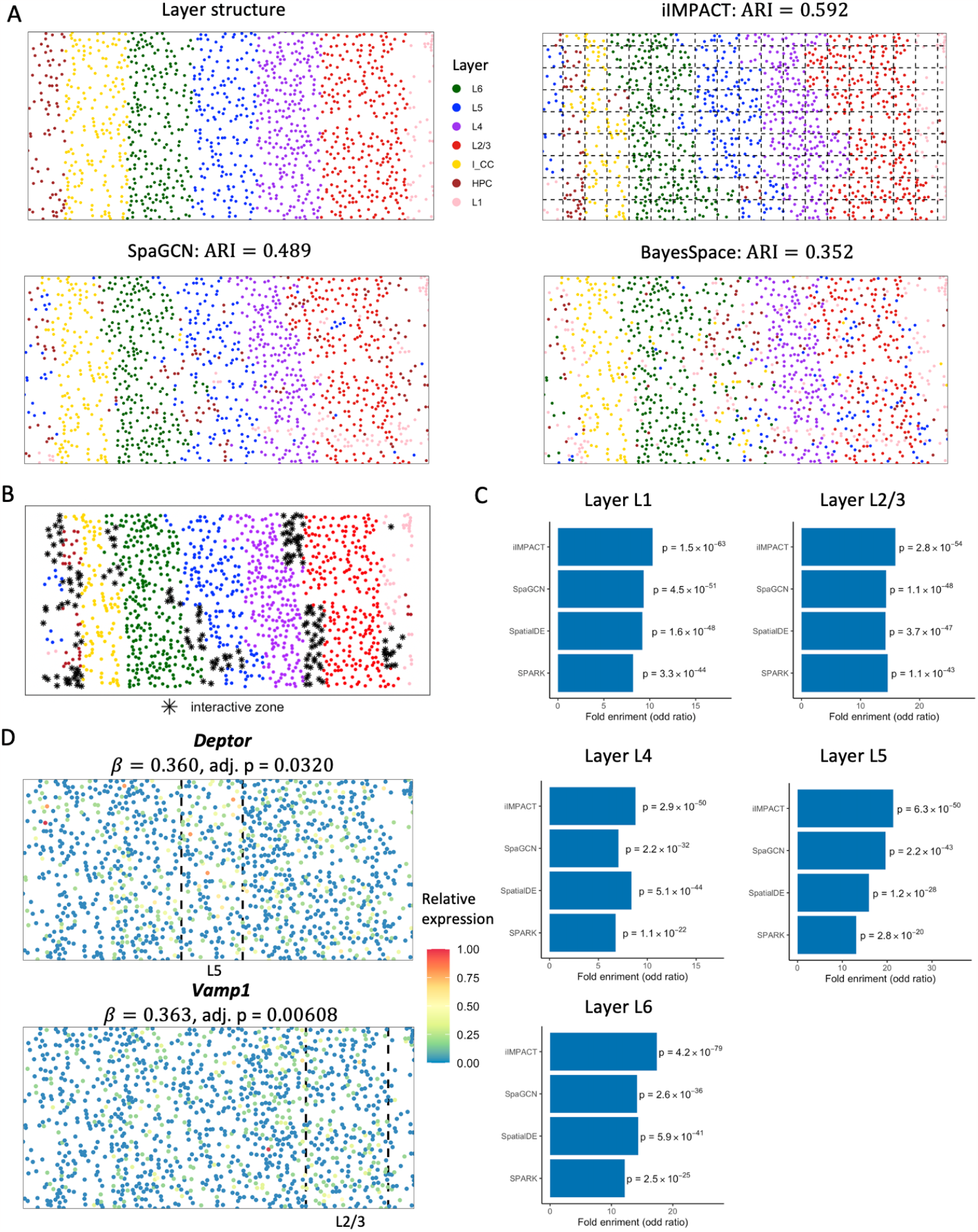
Mouse visual cortex STARmap data: A. Layer structure of the tissue section from the original study. Spatial domains detected by iIMPACT, SpaGCN, and BayesSpace, setting the number of clusters to seven (the number of layers). Manually added square lattice grid when fitting iIMPACT is displayed with dashed lines. B. Interactive zones (black asterisk spots) defined by iIMPACT. C. Gene enrichment analysis between SVGs detected by iIMPACT, SpaGCN, SpatialDE, and SPARK, and genes functionally relevant to visual cortex for five layers. D. Spatial expression patterns of two example SVGs, *Deptor* and *Vamp3*, that were only detected by iIMPACT.

As shown in Figure 5A, iIMPACT displayed the most accurate clustering results with the known layer structure (ARI =0.592). We also noticed that implementing iIMPACT on a lower resolution level (grids in Figure 5A) might reduce the influence of noise, thus making the clustering result more robust. We also leveraged iIMPACT to identify the interactive zones (Figure 5B). The majority areas of identified interacting areas were boundaries between two adjacent layers.

We found these iIMPACT-defined SVGs are frequently functionally relevant to visual cortex (Figure 5C). For example, we observed *Deptor* (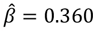, with adjusted p-value = 0.0320), which is highly expressed and functions in a significant portion of corticostriatal and callosal neurons, located in the middle and superficial portions of layer 5 (L5) ^42^, and *Vamp1* (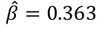, with adjusted p-value = 0.00608), which is ubiquitously expressed and functioned in layer III pyramidal neurons in higher-order areas ^43^ (Figure 5D).

## DISCUSSION

In this paper, we presented iIMPACT, a two-stage statistical method that integrates histology images and molecular profiles. The first stage is a Bayesian finite normal-multinomial mixture model for identifying histology-based spatial domains. Compared with the alternative methods, it fully leverages cellular-level information from histology images to improve clustering performance and increase interpretability. The second stage is a NB regression model for detecting domain-specific SVGs. From both simulation study and real data analysis, we demonstrated that iIMPACT had higher accuracy in identifying spatial domains than published state-of-the-art methods due to the integration of histopathology images in iIMPACT. In addition, iIMPACT is versatile in analyzing both NGS-based and imaging-based SRT techniques, and therefore have broad impacts in the SRT field. Furthermore, iIMPACT has good biological interpretability to characterize histology-based spatial domains. For example, the inferred domain-specific cell-type compositions are consistent with curated annotations, and the interactive zones emphasize the areas with highly heterogeneous cell-type composition and gene expression compared with surroundings. Compared with other SVG detection methods, iIMPACT-defined SVGs are more enriched of known functional genes, confirming that iIMPACT could provide a better understanding of both cellular spatial organization and functional gene landscape of developmental and diseased tissues. Last but not least, compared with other methods, we also confirmed that iIMPACT is computationally efficient (Table S2).

There are several important future extensions for iIMPACT. First, improvement of nuclei segmentation and classification methods might further improve the performance of iIMPACT and therefore will be our focus in the near future. Second, the number of histology-based spatial domains has to be pre-specified when implementing the current version of iIMPACT. To automatically estimate the number of spatial domains, we plan to replace the proposed Bayesian finite mixture model with a Bayesian nonparametric model, such as the Dirichlet process mixture model ^44^ or mixture of finite mixture model ^45, 46^. Third, cell-cell interaction information can be incorporated into iIMPACT to improve the accuracy of histology-based spatial domain identification and increase the model interpretability. These future directions could potentially further boost the performance and interpretability of iIMPACT.

## METHODS

In this section, we first define the molecular and geospatial profiles from NGS-based SRT data (e.g., spatial transcriptomics and the improved 10x Visium platform) and the image profile from the matching AI-reconstructed histology image. Then we discuss how to construct the corresponding profiles from imaging-based SRT (e.g., STARmap) data. After that, we detail the statistical models used in the two stages of iIMPACT. Table S3 in the supplementary material summarizes all key notations introduced in this section.

## Data preparation

### Molecular profile *Y*

In general, the spot-level molecular profile of NGS-based SRT data can be represented by an *N* × *P* count table ***C***, where each entry *c*_*ij*_ ∈ ℕ, *i* = 1, …, *N, j* = 1, …, *P* is the read count for gene *j* measured at spot *i*. To account for nuisance effects across spots, including sequencing depth, amplification and dilution efficiency, and reverse transcription efficiency, we normalize each read count *c*_*ij*_ to its relative level 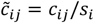, where *s*_*i*_ is the total sum of counts across all genes at spot *i*, 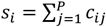, although other normalization methods are acceptable. Then, the relative gene expression 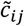 are further log transformed to approximately conform to normality. Following the preprocessing steps in BayesSpace ^18^, we select the top 2,000 most highly variable genes in terms of their relative expression and perform principal component analysis (PCA), or other dimension reduction techniques (e.g., t-SNE ^47^ or UMAP ^48^), to obtain the low-dimensional representation of normalized gene expression denoted by an *N* × *P*′ matrix ***Y***, where each entry *y*_*ij*_ ∈ ℝ, *i* = 1, …, *N, j* = 1, …, *P*′ is the value of the *j*-th top principal component (PC) at spot *i*. We choose to model the PCs in ***Y*** rather than the raw count table ***C*** to avoid the use of complex finite mixture models with feature selection based on cumbersome multivariate distributions. Here, we recommend modeling the top three PCs (*P*′ = 3) for simplicity. A sensitivity analysis on the human breast cancer data (see Figure S6) shows that larger *P*, only provided marginal improvements in clustering performance.

### Image profile *V*

To integrate the image profile into iIMPACT, we applied a nuclei segmentation and identification algorithm, the histology-based digital (HD)-Staining model ^20^, to extract cellular features from images. The HD-Staining model is a trained deep-learning model implemented by the mask regional convolutional neural network (Mask R-CNN) architecture ^49^ for the tumor morphological microenvironment to segment the nuclei of different types of cells, such as immune, tumor and stromal cells. The model was first trained using histology images from lung adenocarcinoma patients in the National Lung Screening Trial study, which has nuclei of six different cell types manually labeled by pathologists. Although the model was originally trained by lung cancer data, it has been improved and verified to be widely adapted to histology image datasets with other cancer types, such as breast cancer, head and neck cancer, ovarian cancer, prostate cancer, and other carcinomas.

The HD-Staining model takes a batch of high-resolution histology image patches of a tissue section as input and simultaneously segments and classifies cell nuclei on this image patch. It provides the locations and types for all identified nuclei in the whole histology image. To match the molecular information measured at spots, which only take less than half area (e.g., the area of all spots in 10x Visium platform is about 38% of the whole domain area), we count cells with different types within each spot and its expanded area (see Figure S7) so that all the cellular information can be utilized. The result is summarized into an *N* × *Q* count matrix ***V***, namely cell abundance table, where each entry *v*_*iq*_ ∈ ℕ, *i* = 1, …, *N, q* = 1, …, *Q* is the number of cells with type *q* observed at spot *i* and its expanded area. iIMPACT leverages the single-cell level histology information from the image profile to enhance spatial domain identification.

### Geospatial profile *G*

Spots are the round area of barcoded mRNA capture probes where gene expression is measured. The spatial distribution of spots is arrayed on a square or triangular lattice. We denote the SRT geospatial profile by an *N* × 2 matrix ***T***, where each row ***t***_*i*_ = (*t*_*i*1_, *t*_*i*2_) gives the *x* and *y* coordinates of the spot *i* on a two-dimensional Cartesian plane. ST and 10x Visium spots are arranged on square and triangular lattice grids, respectively. Thus, defining a neighborhood structure provides an alternative way to represent the geospatial profile ***G***. In particular, ***G*** is an *N* × *N* binary adjacent matrix, where each entry *g*_*ii*,_ = 1 if spot *i* and *i*′ are neighbors (i.e., the Euclidean distance 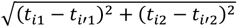 between spot *i* and *i*^′^ is less than a threshold) and *g*_*ii*′_ = 0 otherwise. Note that each diagonal entry *g*_*ii*′_ is equal to zero. There are four and six neighbors for each non-boundary spot from the ST and 10x Visium platforms, respectively. With this neighborhood structure ***G*** as our geospatial profile, the spatial information can be easily integrated into Bayesian cluster analysis via an appropriate prior setting.

### Special handling to imaging-based SRT data

Imaging-based SRT techniques usually have a higher spatial resolution than NGS-based SRT techniques, which is capable of profiling mRNA at the single-cell level. Data from some imaging-based platforms might provide the spatial distribution and types of cells on the tissue section in the original study. To fit iIMPACT to imaging-based SRT data such as STARmap ^11^, we manually add a square lattice grid with appropriate size to the whole domain and consider each square unit as a spot (see Figure 5A). Note that those ‘spots’ fill the whole domain; thus, there is no gap between two adjacent spots. For STARmap data in the RESULTS section, the grid size was chosen to be 750 × 750 pixels, resulting in *N* = 170 spots. Each non-boundary spot has four neighboring spots. We define ***G*** with each entry *g*_*ii*′_ = 1 if spot *i* and *i*′ are neighbors. To construct the molecular profile ***Y***, we first normalize, transform, and reduce the dimension of the gene expression counts at the single-cell level, and then average the resulting values across all cells within each spot. To obtain the “image” profile ***V***, we directly count the cells with different types in each spot.

### Stage I: A Bayesian normal-multinomial mixture model for identifying histology-based spatial domains

The first stage of iIMPACT is to use a Bayesian finite mixture model to partition the whole domain into *K* mutually exclusive histology-based spatial domains. In general, a finite mixture model ^50, 51^ generates random variables from a weighted sum of *K* independent distributions that belong to the same parametric family,

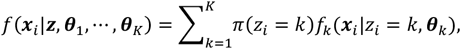

where ***z*** = (*z*_1_, …, *z*_*N*_)^T^ denotes the latent variables specifying the identity of the mixture component *f*_*k*_, characterized by ***θ***_*k*_, to each observation ***x***_*i*_. In the context of this paper, 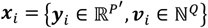 represents the observed molecular and image profiling data, and *z*_*i*_ = *k* indicates that spot *i* belongs to histology-based spatial domain *k*. Since there are two modalities ***Y*** and ***V***, we decompose the mixture component *f*_*k*_ into two sub-components described below. In addition, we incorporate the information from the geospatial profile ***G*** into the prior placed over the auxiliary variable ***z***, encouraging the neighboring spots to be in the same histology-based spatial domain. The number of histology-based spatial domains *K* is decided by prior biological knowledge when available or otherwise by the elbow of the Bayesian information criterion (BIC) plot ^52^.

### Modeling the molecular profile *Y*

We use a multivariate normal (MN) sub-component for modeling the low-dimensional gene expression ***y***_*i*_ at spot *i*:

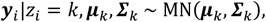

where ***μ***_*k*_ = (*μ*_*k*1_, …, *μ*_*kP′*_)^T^, *μ*_*kP*_ ∈ ℝ is the domain-specific mean vector and ***Σ***_*k*_ is the *P*^′^ × *P*′ domain-specific variance-covariance matrix, requiring positive definiteness. For computational efficiency, we specify a normal prior for ***μ***_0_ conditional on ***Σ***_*k*_, and an inverse-Wishart (IW) prior for ***Σ***_*k*_, i.e., ***μ***_*k*_|***Σ***_*k*_ ∼ M*N*(***ν***_0_, ***Σ***_*k*_/*τ*_0_) and ***Σ***_*k*_ ∼ IW(*η*_0_, ***Φ***_0_). This conjugate setting leads to analytically tractable posterior distributions on ***μ***_*k*_ and ***Σ***_*k*_. Here, ***ν***_0_, *τ*_0_, *η*_0_, and F_0_ are fixed hyperparameters. We set ***ν***_0_ to be the empirical mean vector over all spots and *τ*_0_ = 0.01 to provide a weak prior information so that the data itself would dominate the estimation of ***μ***_*k*_. We set the degree of freedom parameter *η*_0_ = *P*^′^ + 1, controlling the informative strength, and the scale matrix ***Φ***_0_ to be the identity matrix. Let 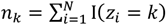 and 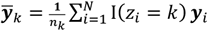 the closed-form posterior distributions are ***μ***_*k*_|***Σ***_*k*_, ***Y*** ∼ M*N*(***ν***_*k*_, ***Σ***_*k*_/*τ*_*k*_) and ***Σ***_*k*_|***Y***∼ IW(*η*_*k*_, ***Φ***_*k*_), where 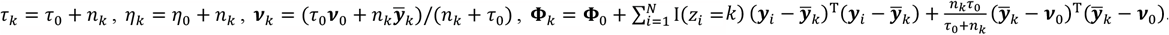

Suppose we choose PCA to perform an orthogonal projection of the scaled and normalized SRT molecular profiling data, we can further set all off-diagonal entries in ***Σ***_*k*_ to be zero, i.e., *σ*_*kpp′*_ = 0, ∀*p* ≠ *p*′. In this case, the multivariate normal model can be decomposed into a product of *P*′ independent normal model,

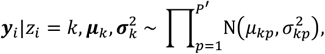

where 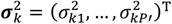 is the set of diagonal entries in ***Σ***_*k*_. The conjugate setting for each dimension becomes a normal-inverse-gamma (IG) distribution ^53^, 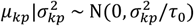 and 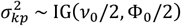, resulting in the closed-form posteriors 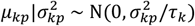 and 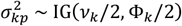, where *τ*_*k*_ = *τ*_0_ + *n*_*k*_, *η*_*k*_ = *η*_0_ + *n*_*k*_, and 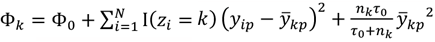. One standard way of setting a weakly informative IG prior is to choose small values of both parameters, such as *ν*_0_/2 = F_0_/2 = 0.1.

### Modeling the image profile *V*

We use a multinomial sub-component for modeling the number of cells with different types ***v***_*i*_ within spot *i* and its expanded area:

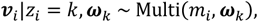

where 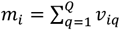 is the total number of cells observed within the area and ***ω***_*k*_ = (*ω*_*k*1_, …, *ω*_*kQ*_)^T^ is defined on a *Q*-dimensional simplex (i.e., *ω*_*kq*_ > 0, ∀*q* and 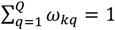 representing the underlying relative abundance of cell types in histology-based spatial domain *k*. Of particular note is that ***ω***_1_, …, ***ω***_*k*_ are the parameters of key interest in iIMPACT, because it can be used to interpret or even define the identified histology-based spatial domains. For example, if a histology-based spatial domain is heavily dominated by cell type *q*, i.e., *ω*_*kq*_ ≫ *ω*_*kq*,_, ∀*q*′, then it could be named after cell type *q*. Note that cell type abundance is assumed to be homogeneous across the same histology-based spatial domain. For computational efficiency, we specify a Dirichlet prior setting for ***ω***_*k*_, i.e., ***ω***_*k*_ ∼ D*i*r(***α***_0_), where ***α***_0_= (*α*_01_, …, *α*_0*Q*_)^T^, *α*_0*q*_ ∈ ℝ^+^ are fixed hyperparameters. This conjugate setting leads to an analytically tractable posterior distribution on ***ω***_*k*_|***V*** ∼ D*i*r*ic*hlet(***α***_*k*_) with each entry 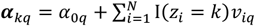. We recommend *α*_01_ = … = *α*_0*Q*_ = 1/2 or 1 for a non or weakly informative setting.

### Incorporating the geospatial profile *G*

To utilize the available spatial information in the geospatial profile, we employ a Markov random field prior ^30, 31^ on the histology-based spatial domain indicator ***z***, encouraging neighboring spots to be clustered into the same histology-based spatial domain:

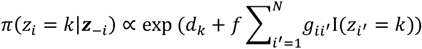

where ***z***_−*i*_ denotes the set of all entries in ***z*** excluding the *i* -th one, the hyperparameters ***d*** = (***d***_1_, …, ***d***_*N*_)^T^ control the number of spots belonging to each of the *K* histology-based spatial domains and *f* ∈ ℝ^+^ controls the spatial dependency or smoothness. Note that if a spot has no neighbors, the above prior distribution reduces to a multinomial distribution, 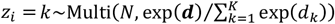. Although the larger the *f*, the smoother the pattern of spatial domains, careful determination of *f* is required. This is because a large value of *f* may lead to a phase transition problem (i.e., all spots are assigned to the same histology-based spatial domain). In this paper, we choose ***d***_1_ = … = ***d***_*k*_ = 1 and *f* = 1 by default, as this setting performs very well in the simulation study and yields reasonable results in our real data analysis.

### Posterior sampling *via* MCMC algorithm

iIMPACT integrates the molecular, image, and geospatial profiles to partition the whole domain into *K* biologically meaningful spatial domains. Because the low-dimensional molecular profile ***Y*** and AI-reconstructed image profile ***V*** are generated from different sources, they are conditionally independent of each other. Thus, we define the mixture component

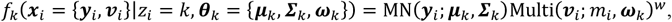

where the tuning parameter ***w*** ∈ [0,1] controls the image profile’s contribution to the clustering process, with respect to that of the molecular profile. Parameterizing the data likelihood above by decreasing ***w*** will result in a flatter multinomial distribution, thus downplaying the role of the image profile. When ***w*** = 0, iIMPACT will not depend on any cell type abundance information. We conducted a sensitivity analysis to search for the best choice of ***w***. Our result suggests setting ***w*** = 0.05 and 0.5 for 10x Visium and STARmap data, respectively (see Figure S8). Note that in addition to the SRT platform and application, we should also consider the image and molecular profiles’ dimensionalities (i.e., *Q* and *P*’) to determine the value of ***w*** with some degree of caution. Finally, we give the full posterior distribution as,

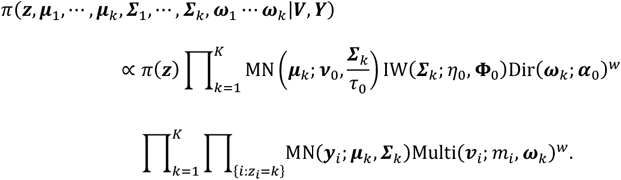

To identify histology-based spatial domains, the posterior distribution of *z*_*i*_ will be of direct interest to us, given by

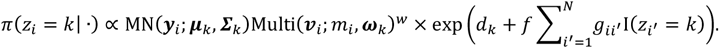

The individual quantities of all possible values of *z*_*i*_ are first computed and then summed to find the normalization constant 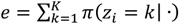. A new value of *z*_*i*_ can be drawn from a multinomial distribution Mult*i*(1, (*π*(*z*_*i*_ = *k*| ·)/*e*, …, *π*(*z*_*i*_ = *K*| ·)/*e*)^T^). For any particular domain-specific parameters, i.e., ***μ***_*k*_, ***Σ***_*k*_, ***ω***_*k*_, we only require the partial data likelihood in estimating its posterior density as detailed before. Since the posterior conditional distributions for all parameters are in closed form, it is straightforward to use a Gibbs sampler, a type of Markov chain Monte Carlo (MCMC) algorithm, to obtain a sequence of observations approximated from the multivariate distribution *π*(***z, μ***_1_, …, ***μ***_*k*_, ***Σ***_1_, …, ***Σ***_*k*_, ***ω***_1_ … ***ω***_*k*_|***V, Y***). Consequently, the posterior inference can be made by post-processing the MCMC samples, such as 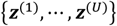 and 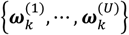, where *u* indexes the MCMC iteration and *U* is the total number of iterations after burn-in.

In any finite mixture model, the invariance of the likelihood under permutation of the cluster labels ***z*** may result in an identifiability problem, leading to symmetric and multimodal posterior distributions with up to *K*! copies of each genuine model. What is worse, it will also complicate inference on other parameters. To address this issue, we impose an order restriction on the posterior samples of parameters ***ω***_1_ … ***ω***_*k*_ based on a given cell type *q*. In particular, at each iteration *u*, we relabel ***z*** and switch all the related domain-specific parameters of the MCMC outputs to satisfy the constraint 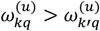 for cluster indicator *k* < *k*′. In other words, the first histology-based spatial domain has the largest proportion of cell type *q*, while histology-based spatial domain *K* has the small proportion of cell type *q*.

### Identifying histology-based spatial domains and interactive zones

Our primary interest lies in identifying histology-based spatial domains via making inferences on the spatial domain indicator vector ***z***. Here we apply the mode estimates ^54^ based on the marginal probabilities 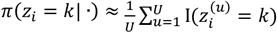. The estimate of 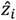 can be obtained by selecting the highest value:

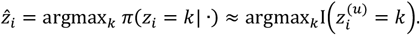

Uncertainty quantification is one advantage of the proposed Bayesian finite mixture model. For example, if the marginal probability of assigning spot *i* to histology-based spatial domain *k* is considerably high, e.g., *π*(*z*_*i*_ = *k*| ·) ≥ 0.9, then we are confident about the assignment. However, if some marginal probabilities are almost equivalent or there is no significant mode for a spot, e.g., *π*(*z*_*i*_ = *k*|) < 0.9, ∀*k*, then we tend not to assign the spot to any histology-based spatial domains. Instead, we define the spot as the boundary spot, and the resulting connected area as the interactive zone.

### Interpreting and defining histology-based spatial domains

The domain-specific relative abundance of cell types ***ω***_1_, …, ***ω***_*k*_ are another group of parameters of interest in our model, because it can be used to interpret or even define the identified histology-based spatial domains. We use the posterior mean as the estimate,

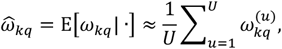

averaging over all its MCMC samples. Additionally, the credible interval for each *ω*_*kq*_ can be approximated by its post-burn-in MCMC sample quantiles. Note that the MCMC samples can also be used to approximate any other quantity of interest that analytical solution is impossible, e.g., *π*(*ω*_*kq*_ > *ω*_*k′ q*_ |) for some *k, k*’, and *q*.

### Stage II: A generalized linear regression model for detecting domain-specific SVGs

To test if each gene is differentially expressed among those identified histology-based spatial domains in Stage I of iIMPACT, we use a generalized linear regression model, where the response variable is gene expression counts, and the predictor variables are the histology-based spatial domain indicators. In particular, we assume that all read counts from a gene *j* across different spots indexed by *i* are from an NB distribution:

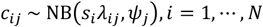

where *s*_*i*_ is the size factor of spot *i*, ψ_)_ is the over-dispersion parameter of gene *j*, and *λ*_*ij*_ is the underlying normalized expression level for gene *j* at spot *i*. We further use the canonical link,

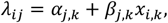

which is typically used in the Poisson and NB regression models. Here, *x*_*i, k*_ = I(*z*_*i*_ = *k*) is a binary indicator. If spot *i* is assigned to histology-based spatial domain *k* in Stage I of iIMPACT, then *x*_*i, k*_ = 1; otherwise, *x*_*i, k*_ = 0. Thus, we can interpret the intercept *α*_*j,k*_ as the baseline expression level of gene *j* in the whole domain excluding histology-based spatial domain *k*, and the slope *β*_*j,k*_ as the differential expression level of gene *j* in histology-based spatial domain *k* as a shift from the baseline. With this modeling framework, SVGs, which are differentially expressed in a given histology-based spatial domain *k* compared with all other domains, can be identified *via* testing the null hypothesis *H*_0_: *β*_*j, k*_ = 0 versus the alternative *H*_α_: *β*_*j, k*_ ≠ 0. For those genes whose resulting adjusted p-values are less than a significance level (e.g., 0.05), we define them as domain-*k*-specific spatially variable genes. To control the false discovery rate, the Benjamini and Hochberg method ^55^ needs to be applied to adjust p-values. The above NB regression model is fitted via the function *glm*.*nb* in the R package MASS ^56^.

## Data availability

The authors analyzed four publicly available SRT datasets. Raw count matrices, images, and spatial data for three SRT datasets from 10x Visium are accessible on the 10x Genomics website at https://support.10xgenomics.com/spatial-gene-expression/datasets. Mouse visual cortex STARmap data can be downloaded from https://www.starmapresources.com/data.

## Code availability

An open-source implementation of the iIMPACT algorithm in R/C++ is available at https://github.com/Xijiang1997/iIMPACT.

## Acknowledgements

This work was supported by the following funding: the National Science Foundation [2210912, 2113674] and the National Institutes of Health [1R01GM141519] (to Q. L.); the National Institutes of Health [R01GM140012, R01GM141519, R01DE030656, U01CA249245], and the Cancer Prevention and Research Institute of Texas [CPRIT RP230330] (to G. X.); the Rally Foundation, Children’s Cancer Fund (Dallas), the Cancer Prevention and Research Institute of Texas (RP180319, RP200103, RP220032, RP170152 and RP180805), and the National Institutes of Health (R01DK127037, R01CA263079, R21CA259771, UM1HG011996, and R01HL144969) (to L. X.); The funding bodies had no role in the design, collection, analysis, or interpretation of data in this study.

