## Supplementary notes for "Integrating Image and Molecular Profiles for Spatial Transcriptomics Analysis"

[2]Department of Statistical Science,
Southern Methodist University,
Dallas, TX 75275, United States.

[3]Department of Pathology,
The University of Texas Southwestern Medical Center,
Dallas, TX 75390, United States.

[4]Department of Mathematical Sciences,
The University of Texas at Dallas,
Richardson, TX 75080, United States.

### S1. Details of the MCMC Algorithm

According to the model description in the METHODS section, the full data likelihood of the proposed Bayesian finite normal-multinomial mixture model is given as follows.

$$\begin{aligned}
& f(\boldsymbol{Y}, \boldsymbol{V} | \boldsymbol{z}, \boldsymbol{\mu}_1, \ldots, \boldsymbol{\mu}_K, \boldsymbol{\Sigma}_1, \ldots, \boldsymbol{\Sigma}_K, \boldsymbol{\omega}_1, \ldots, \boldsymbol{\omega}_K) \\
=& \prod_{k=1}^{K} \prod_{i=1}^{N} \mathrm{I}(z_i = k) f(\boldsymbol{y}_i, \boldsymbol{v}_i | z_i = k, \boldsymbol{\mu}_k, \boldsymbol{\Sigma}_k, \boldsymbol{\omega}_k),
\end{aligned}$$

where $\mathrm{I}(\cdot)$ denotes the indicator function and

$$\begin{aligned}
& f(\boldsymbol{y}_i, \boldsymbol{v}_i | z_i = k, \boldsymbol{\mu}_k, \boldsymbol{\Sigma}_k, \boldsymbol{\omega}_k) \\
&= f(\boldsymbol{y}_i | z_i = k, \boldsymbol{\mu}_k, \boldsymbol{\Sigma}_k) f(\boldsymbol{v}_i | z_i = k, \boldsymbol{\omega}_k)^w \\
&= \mathrm{MN}(\boldsymbol{y}_i; \boldsymbol{\mu}_k, \boldsymbol{\Sigma}_k) \mathrm{Multi}(\boldsymbol{v}_i; m_i, \boldsymbol{\omega}_k)^w \\
&= (2\pi)^{-P'/2} |\boldsymbol{\Sigma}_k|^{-1/2} \exp\left(-\frac{1}{2}(\boldsymbol{y}_i - \boldsymbol{\mu}_k)^\top \boldsymbol{\Sigma}_k^{-1} (\boldsymbol{y}_i - \boldsymbol{\mu}_k)\right) \left(\frac{m_i!}{\prod_{q=1}^{Q} v_{iq}!} \prod_{q=1}^{Q} \omega_{kq}^{v_{iq}}\right)^w \\
&\propto |\boldsymbol{\Sigma}_k|^{-1/2} \exp\left(-\frac{1}{2}(\boldsymbol{y}_i - \boldsymbol{\mu}_k)^\top \boldsymbol{\Sigma}_k^{-1} (\boldsymbol{y}_i - \boldsymbol{\mu}_k)\right) \left(\prod_{q=1}^{Q} \omega_{kq}^{v_{iq}}\right)^w .
\end{aligned}$$

We assume an independent prior structure 1) between the normal and multinomial sub-components; and 2) among their parameters belonging to different spatial domains. Thus, the joint distribution of priors for parameters is given as follows.

$$\begin{aligned}
& \pi(\boldsymbol{z}, \boldsymbol{\mu}_1, \ldots, \boldsymbol{\mu}_K, \boldsymbol{\Sigma}_1, \ldots, \boldsymbol{\Sigma}_K, \boldsymbol{\omega}_1, \ldots, \boldsymbol{\omega}_K) \\
&= \pi(\boldsymbol{z}) \pi(\boldsymbol{\mu}_1, \ldots, \boldsymbol{\mu}_K, \boldsymbol{\Sigma}_1, \ldots, \boldsymbol{\Sigma}_K) \pi(\boldsymbol{\omega}_1, \ldots, \boldsymbol{\omega}_K)^w \\
&= \pi(\boldsymbol{z}) \prod_{k=1}^{K} \pi(\boldsymbol{\mu}_k, \boldsymbol{\Sigma}_k) \prod_{k=1}^{K} \pi(\boldsymbol{\omega}_k)^w \\
&= \pi(\boldsymbol{z}) \prod_{k=1}^{K} \pi(\boldsymbol{\mu}_k | \boldsymbol{\Sigma}_k) \pi(\boldsymbol{\Sigma}_k) \prod_{k=1}^{K} \pi(\boldsymbol{\omega}_k)^w .
\end{aligned}$$

We assign the Markov random field (MRF) prior for histology-based spatial domain indicator

$\boldsymbol{z}$ as

$$\pi(z_i|\boldsymbol{z}_{-i}) \propto \exp\left(d_k + f\sum_{i'=1,i'\neq i}^{N} g_{ii'}\mathrm{I}(z_{i'}=k)\right),$$

We assign the conjugate priors for other parameters, listed as follows, so that the Gibbs sampler can be applied for posterior sampling.

$$\boldsymbol{\mu}_k|\boldsymbol{\Sigma}_k \sim \mathrm{MN}(\boldsymbol{\nu}_0, \boldsymbol{\Sigma}/\tau_0)$$
$$\text{or equivalently, } \pi(\boldsymbol{\mu}_k|\boldsymbol{\Sigma}_k) = (2\pi/\tau_0)^{-P'/2}|\boldsymbol{\Phi}_0|^{-1/2}\exp\left(-\frac{\tau_0}{2}(\boldsymbol{\mu}_k-\boldsymbol{\nu}_0)^\top\boldsymbol{\Phi}_0^{-1}(\boldsymbol{\mu}_k-\boldsymbol{\nu}_0)\right),$$
$$\boldsymbol{\Sigma}_k \sim \mathrm{IW}(\eta_0, \boldsymbol{\Phi}_0)$$
$$\text{or equivalently, } \pi(\boldsymbol{\Sigma}_k) = \frac{|\boldsymbol{\Phi}_0|^{\eta_0/2}}{2^{\eta_0 P'/2}\Gamma_{P'}(\eta_0/2)}|\boldsymbol{\Sigma}_k|^{-(\eta_0+P'+1)/2}\exp\left(-\frac{1}{2}\mathrm{tr}\left(\boldsymbol{\Phi}_0\boldsymbol{\Sigma}_k^{-1}\right)\right),$$

and

$$\boldsymbol{\omega}_k \sim \mathrm{Dir}(\boldsymbol{\alpha}_0) \text{ or equivalently, } \pi(\boldsymbol{\omega}_k) = \frac{\Gamma\left(\sum_{q=1}^{Q}\alpha_{0q}\right)}{\prod_{q=1}^{Q}\Gamma(\alpha_{0q})}\prod_{q=1}^{Q}\omega_{kq}^{\alpha_{0q}-1},$$

where $\Gamma_{P'}(\cdot)$ and $\Gamma(\cdot)$ denote the $P'$-dimensional and univariate gamma function.

We recommend a weakly informative prior setting by choosing the MRF hyperparameters $d_1 = \ldots = d_K = 1$ and $f = 1$, the multivariate normal hyperparameters $\boldsymbol{\nu}_0 = \frac{1}{N}\sum_{i=1}^{N}\boldsymbol{y}_i$, $\tau_0 = 0.01$, $\eta_0 = P' + 1$, and $\boldsymbol{\Phi}_0 = \boldsymbol{I}_{P'\times P'}$ (i.e., the $P'$-by-$P'$ identity matrix), and the multinomial hyperparameters $\alpha_{01} = \ldots = \alpha_{0Q} = 1$.

The full posterior distribution of the proposed Bayesian normal-multinomial mixture model is given in the following formula.

$$\pi(\boldsymbol{z}, \boldsymbol{\mu}_1, \ldots, \boldsymbol{\mu}_K, \boldsymbol{\Sigma}_1, \ldots, \boldsymbol{\Sigma}_K, \boldsymbol{\omega}_1, \ldots, \boldsymbol{\omega}_K|\boldsymbol{Y}, \boldsymbol{V}) \propto$$
$$f(\boldsymbol{Y}, \boldsymbol{V}|\boldsymbol{z}, \boldsymbol{\mu}_1, \ldots, \boldsymbol{\mu}_K, \boldsymbol{\Sigma}_1, \ldots, \boldsymbol{\Sigma}_K, \boldsymbol{\omega}_1, \ldots, \boldsymbol{\omega}_K)\pi(\boldsymbol{z}, \boldsymbol{\mu}_1, \ldots, \boldsymbol{\mu}_K, \boldsymbol{\Sigma}_1, \ldots, \boldsymbol{\Sigma}_K, \boldsymbol{\omega}_1, \ldots, \boldsymbol{\omega}_K)$$

Posterior sampling is employed by the MCMC algorithm. Our primary interest lies in identifying histology-based spatial domains and the interactive zone *via* inferring the spatial domain indicator vector $\boldsymbol{z}$, and in characterizing domain-specific relative abundance of cell types *via* inferring $\boldsymbol{\omega}_1, \ldots, \boldsymbol{\omega}_K$. Since we use conjugate priors on all model parameters

$\{\boldsymbol{z}, \boldsymbol{\mu}_1, \ldots, \boldsymbol{\mu}_K, \boldsymbol{\Sigma}_1, \ldots, \boldsymbol{\Sigma}_K, \boldsymbol{\omega}_1, \ldots, \boldsymbol{\omega}_K\}$, their conditional distributions are all in closed form and easy to sample from. Consequently, we can rely on the Gibbs sampler, an MCMC algorithm for obtaining a sequence of observations approximated from a multivariate probability distribution when direct sampling is difficult. To be specific, we perform the following steps sequentially at each MCMC iteration after a random initialization.

**Update the histology-based spatial domain indicator $\boldsymbol{z}$**: We update $z_1, \ldots, z_N$ sequentially. To allocate spot $i$ to one of the $K$ histology-based spatial domains, we sample $z_i$ from a single-drawing multinomial distribution,

$$z_i|\cdot \sim \text{Multi}(1, (\pi(z_i = 1|\cdot)/e, \ldots, \pi(z_i = K|\cdot)/e)),$$

where

$$\begin{aligned}\pi(z_i = k|\cdot) &\propto f(\boldsymbol{y}_i, \boldsymbol{v}_i | z_i = k, \boldsymbol{\mu}_k, \boldsymbol{\Sigma}_k, \boldsymbol{\omega}_k)\pi(z_i = k|\boldsymbol{z}_{-i}) \\ &\propto |\boldsymbol{\Sigma}_k|^{-1/2} \exp\left(-\frac{1}{2}(\boldsymbol{y}_i - \boldsymbol{\mu}_k)^\top \boldsymbol{\Sigma}_k^{-1}(\boldsymbol{y}_i - \boldsymbol{\mu}_k)\right)\left(\prod_{q=1}^{Q} \omega_{k,q}^{v_{iq}}\right)^w \\ &\quad \exp\left(d_k + f \sum_{i'=1, i' \neq i}^{N} g_{ii'} \mathrm{I}(z_{i'} = k)\right)\end{aligned}$$

and the normalization constant $e = \sum_{k=1}^{K} \pi(z_i = k|\cdot)$.

**Update the domain-specific relative abundance of cell types $\boldsymbol{\omega}_k$'s**: We update $\boldsymbol{\omega}_1, \ldots, \boldsymbol{\omega}_K$ sequentially. For each histology-based spatial domain $k$, we draw a sample of $\boldsymbol{\omega}_k$ from a Dirichlet distribution,

$$\boldsymbol{\omega}_k|\cdot \sim \text{Dir}(\boldsymbol{\alpha}_k),$$

where the concentration parameters $\boldsymbol{\alpha}_k = (\alpha_{k1}, \ldots, \alpha_{kQ})$ with each entry $\alpha_{kq} = \alpha_{0q} + \sum_{i=1}^{N} \mathrm{I}(z_i = k) v_{iq}$. Note that the last term $\sum_{i=1}^{N} \mathrm{I}(z_i = k) v_{iq}$ denotes the total number of cells with type $q$ observed in histology-based spatial domain $k$.

**Update the domain-specific low-dimensional representation of gene expression mean $\boldsymbol{\mu}_k$'s**: We update $\boldsymbol{\mu}_1, \ldots, \boldsymbol{\mu}_K$ sequentially. For each histology-based spatial domain $k$, we draw a sample of $\boldsymbol{\mu}_k$ from a multivariate normal distribution,

$$\boldsymbol{\mu}_k|\cdot \sim \mathrm{MN}(\boldsymbol{\nu}_k, \boldsymbol{\Sigma}_k/\tau_k),$$

where $\boldsymbol{\nu}_k = (\boldsymbol{\nu}_0\tau_0 + n_k\bar{\boldsymbol{y}}_k)/(\tau_0 + n_k)$ and $\tau_k = \tau_0 + n_k$. Note that $n_k = \sum_i^N \mathrm{I}(z_i = k)$ is the number of spots allocated to histology-based spatial domain $k$ and $\bar{\boldsymbol{y}}_k = \frac{1}{n_k}\sum_i^N \mathrm{I}(z_i = k)\boldsymbol{y}_i$ denotes the average low-dimensional gene expression value over all the spots allocated to histology-based spatial domain $k$.

If PCA is chosen to reduce the dimension of the SRT molecular profile, then we can further set $\boldsymbol{\Sigma}_k$ to an $P'$-by-$P'$ diagonal matrix due to orthogonality among principal components. In this special case, we can draw each entry in $\boldsymbol{\mu}_k$ independently,

$$\mu_{kj}|\cdot \sim \mathrm{N}(\nu_{kj}, \sigma_{kj}^2/\tau_k),$$

where $\nu_{kj} = (\nu_{0j}\tau_0 + n_k\bar{y}_{kj})/(\tau_0 + n_k)$.

**Update the domain-specific covariance matrix of the low-dimensional representation of gene expression $\boldsymbol{\Sigma}_k$'s**: We update $\boldsymbol{\Sigma}_1, \ldots, \boldsymbol{\Sigma}_K$ sequentially. For each histology-based spatial domain $k$, we draw a sample of $\boldsymbol{\Sigma}_k$ from an inverse-Wishart distribution,

$$\boldsymbol{\Sigma}_k|\cdot \sim \mathrm{IW}(\eta_k, \boldsymbol{\Phi}_k),$$

where $\eta_k = \eta_0 + n_k$ and $\boldsymbol{\Phi}_k = \boldsymbol{\Phi}_0 + \sum_{i=1}^N \mathrm{I}(z_i = k)(\boldsymbol{y}_i - \bar{\boldsymbol{y}}_k)(\boldsymbol{y}_i - \bar{\boldsymbol{y}}_k)^\top + \frac{\tau_0 n_k}{\tau_0 + n_k}(\bar{\boldsymbol{y}}_k - \boldsymbol{\nu}_0)(\bar{\boldsymbol{y}}_k - \boldsymbol{\nu}_0)^\top$.

For the case when using PCA, the inverse-Wishart prior reduces to an inverse-gamma prior, $\sigma_{kj}^2 \sim \mathrm{IG}(\eta_0/2, \phi_0/2)$. Thus, we only need to draw each diagonal entry in $\boldsymbol{\Sigma}_k$ independently,

$$\sigma_{kj}^2|\cdot \sim \mathrm{IG}(\eta_k/2, \phi_k/2),$$

where $\phi_k = \phi_0 + \sum_{i=1}^N \mathrm{I}(z_i = k)(y_{ij} - \bar{y}_{kj})^2 + \frac{\tau_0 n_k}{\tau_0 + n_k}(\bar{y}_{kj} - \nu_{0j})^2$.

### S2. Simulation Study

**Data generative model:** The simulated data were generated based on the $K = 5$ histology-based spatial domains identified by iIMPACT on the human breast cancer dataset (Figure 3B). The posterior means of those domain-specific relative abundances of cell types $\hat{\boldsymbol{\omega}}_k$'s and mean vectors of low-dimensional representation of gene expression $\hat{\boldsymbol{\mu}}_k$'s are given in Table S4. Using this real data information, we generated the cell type abundance for each spot from a multinomial distribution

$$\boldsymbol{v}_i | z_i = k \;\sim\; \text{Multi}(m_i, \hat{\boldsymbol{\omega}}_k),$$

where the size parameter $m_i$ was also obtained from real data. For generating high-dimensional gene expression counts, we first projected the $P'$-dimensional domain-specific mean vectors $\hat{\boldsymbol{\mu}}_k$'s on to the original basis, denoted by a $P$-dimensional vector $\tilde{\boldsymbol{\mu}}_k$. To mimic the excess zeros and over-dispersion, we sampled each gene expression count $c_{ij}$ from a zero-inflated negative binomial (ZINB) distribution,

$$c_{ij} | z_i = k \;\sim\; \pi_i \text{I}(c_{ij} = 0) + (1 - \pi_i)\text{NB}(s_i \exp(\tilde{\mu}_{kj}), \psi_j), \tag{1}$$

where the size factors $s_i$'s were sampled from a log-normal distribution with mean 0 and standard deviation 0.2, i.e., $s_i \sim \text{LN}(0, 0.2)$. The dispersion parameters $\psi_j$'s were sampled from an exponential distribution, i.e., $\psi_j \sim \text{Exp}(\lambda_\psi)$, with two choices of the rate parameter $\lambda_\psi = 0.1$ or $0.2$, corresponding to low and high variability. The false zero proportion parameters $\pi_i$'s were set to be 0.3 or 0.5, corresponding to low and high sparsity. For each of the four scenarios in terms of $\lambda_\psi$ and $\pi_i$, we independently repeated the above steps to generate 10 replicated simulated datasets.

**iIMPACT settings:** We chose the number of reduced dimensions as $P' = 3$ in the PCA step for obtaining the low-dimensional representation of gene expression levels $\boldsymbol{Y}$. The number of histology-based spatial domains was fixed at $K = 5$. We followed the recommended prior setting, as detailed in the METHODS section. As for the MCMC algorithm, we ran four independent MCMC chains with $U = 10,000$ iterations, discarding the first half as burn-in. We started each chain from a model by randomly drawing all parameters from their prior distributions. The results reported in Figure S9 were obtained by pooling the MCMC outputs from the four chains after.

**Competing methods:** We compared the performance of iIMPACT on spatial domain identification with four current state-of-the-art methods, Louvain (Blondel et al., 2008; Satija et al., 2015), stLearn (Pham et al., 2020), BayesSpace (Zhao et al., 2021), and SpaGCN (Hu et al., 2021). We used the default setting of each competing method, as suggested by the authors. The number of spatial domains was fixed as $K = 5$ for all methods.

**Results:** We quantified the clustering performance *via* the widely used adjusted Rand index (ARI). It ranges from $-1$ to 1, with higher values indicating greater consistency between the clustering results and the ground truth. The results are shown in Figure S9. iIMPACT substantially outperformed all other methods, exhibiting the highest average ARI, under all four scenarios, which highlights the benefit of integrating the cell type abundance information into the spatial domain identification process. SpaGCN also demonstrated superior performance compared to other competing methods, leveraging its ability to effectively utilize histology information. Conversely, stLearn, despite its capability to incorporate histology images, had unsatisfactory clustering accuracy and performed similarly to Louvain, a non-spatial clustering method. BayesSpace had a large variance among replicates since it might fail to converge for some replicates. Comparing the performance under low and high variability settings, iIMPACT was robust to the level of over-dispersion of gene expression counts due to the normalization and dimensionality reduction procedures employed before the clustering model. However, it suffered from reduced clustering accuracy under high sparsity settings.

### S3. Sensitivity Analysis

We first conducted a sensitivity analysis to investigate how the number of dimensions reduced affects iIMPACT's performance in histology-based spatial domain identification. In particular, we varied the number of top principal components (PC) $P'$ kept in the molecular profile from 2 to 15 and computed the ARI between the ground truth and the MAP estimate of the histology-based spatial domain indicator $\hat{\boldsymbol{z}}$. Take human breast cancer data as example, Figure S6 summarizes the achieved ARI against different choices of $P'$, along with the cumulative variance explained by the PCs, indicating that iIMPACT was relatively robust to the choice of number of PCs and the best performance occured when $P' = 3$.

Then, another sensitivity analysis was performed to investigate the sensitivity of iIMPACT to the choice of the image profile weight $w$, which controls the image profile's contribution to the spatial domain identification result. In particular, we varies $w$ in

$$\{0.01, 0.02, 0.05, 0.10, 0.25, 0.50, 0.75, 1.00\}$$

and computed the ARI between the ground truth and the MAP estimate of the histology-based spatial domain indicator $\hat{\boldsymbol{z}}$. Figure S8 summarizes the achieved ARI against different values of $w$ for both the human breast cancer and mouse visual cortex STARmap data. It is interesting to know that iIMPACT was very sensitive to the choice of $w$. The best performance occured when setting $w = 0.05$ for the human breast cancer data from NGS-based SRT platforms (e.g., 10x Visium) and $w = 0.5$ for the mouse visual cortex data from imaging-based SRT techniques (e.g., STARmap). Therefore, to guard against under or over-fitting, the value of $w$ should be chosen with some degree of caution.

### REFERENCES

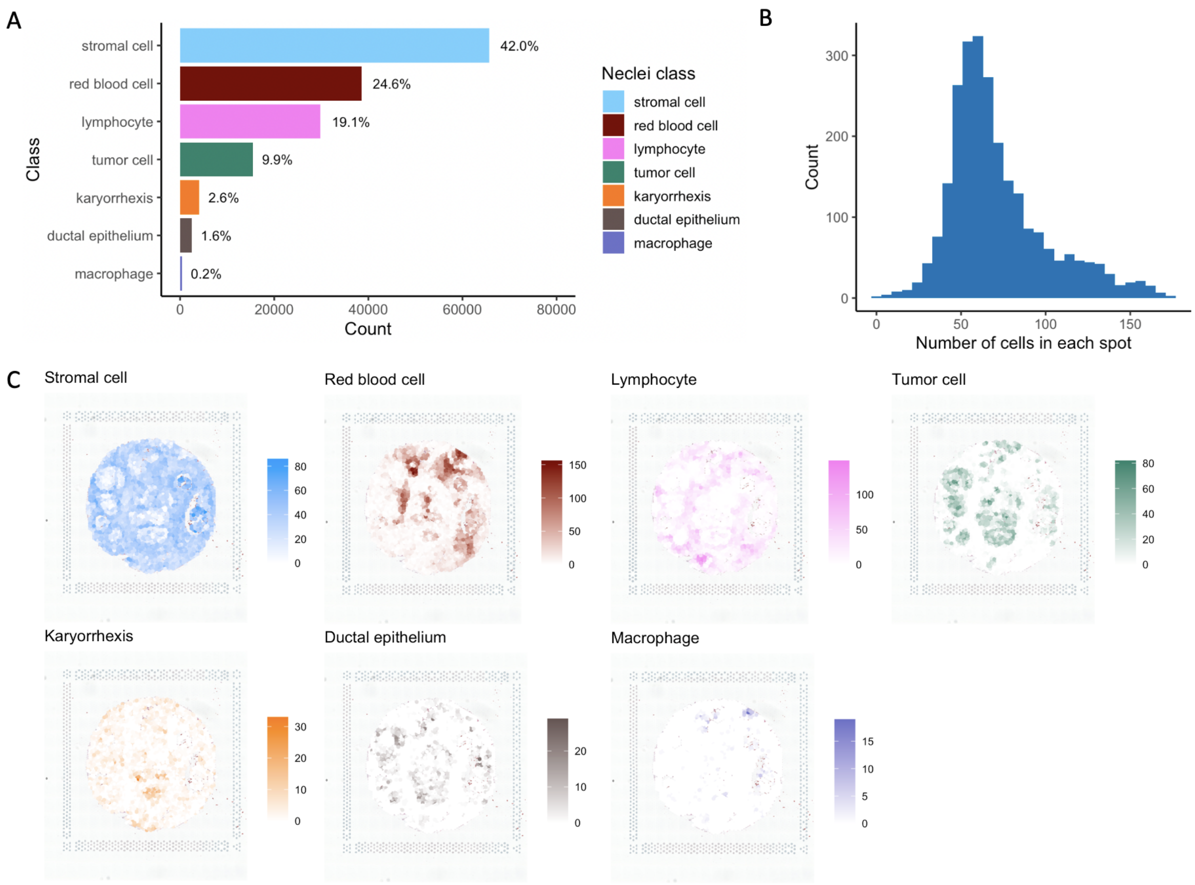


**Figure S1**: Nuclei identification results from the HD-Staining model for human breast cancer data: A. The number of nuclei identified for seven different nuclei classes; B. Histogram of number of cells in each spot expanded area; C. Spatial distribution of spot-level cell type abundance for seven nuclei classes.

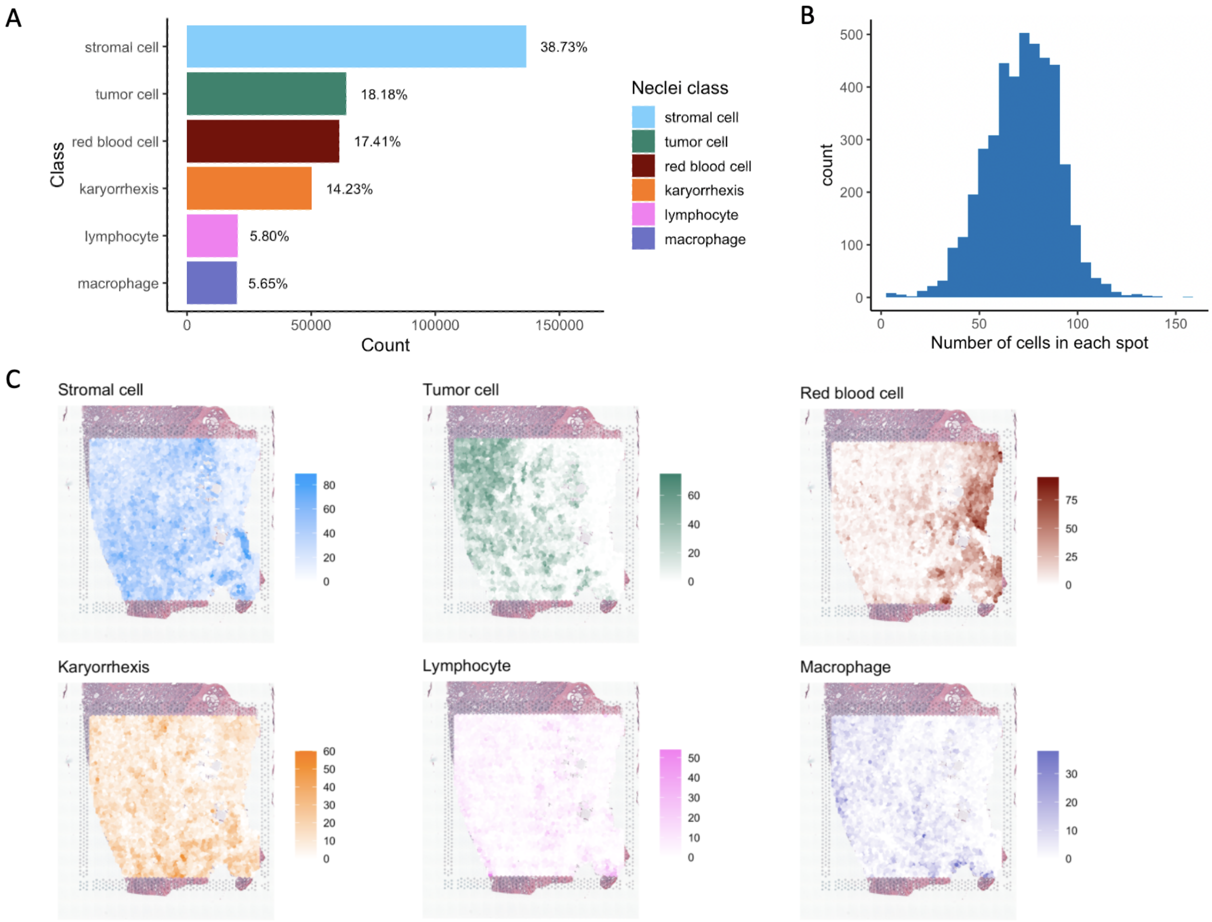


**Figure S2**: Nuclei identification results from the HD-Staining model for human prostate cancer data: A. The number of nuclei identified for six different nuclei classes; B. Histogram of number of cells in each spot expanded area; C. Spatial distribution of spot-level cell type abundance for six nuclei classes.

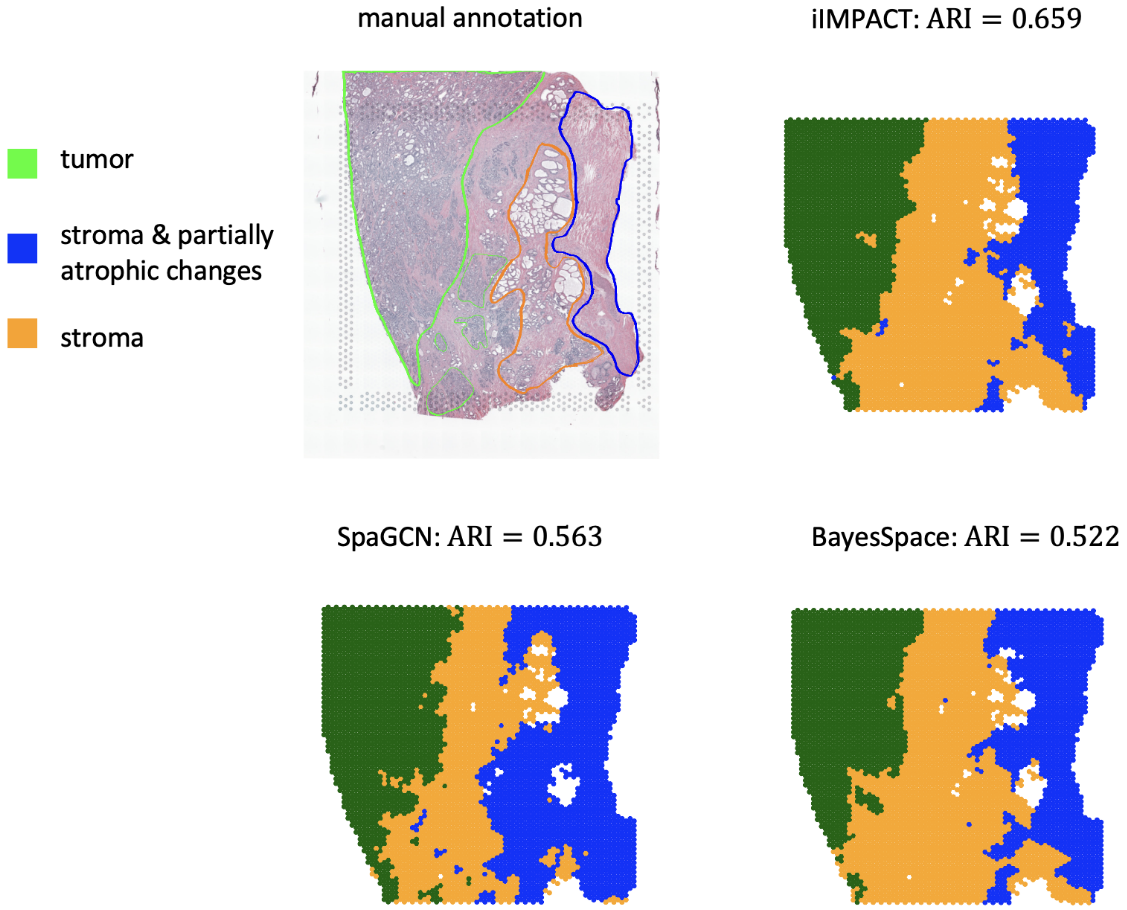


**Figure S3**: Histology image of the tissue section with manually annotated labels from pathologists, and histology-based spatial domains detected by iIMPACT, SpaGCN, and BayesSpace, setting the number of clusters to be 3, for human prostate cancer data.

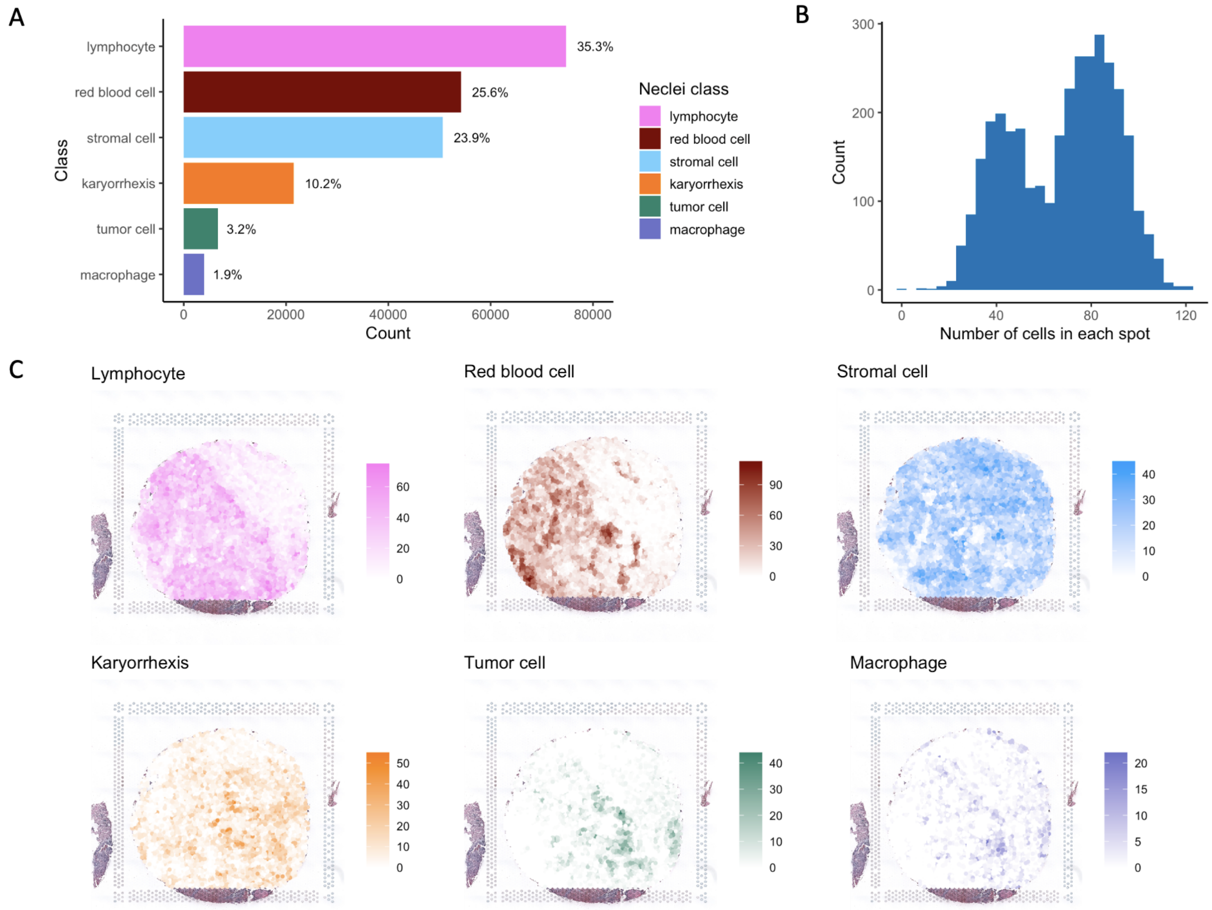


**Figure S4**: Nuclei identification results from the HD-Staining model for human ovarian cancer data: A. The number of nuclei identified for six different nuclei classes; B. Histogram of number of cells in each spot expanded area; C. Spatial distribution of spot-level cell type abundance for six nuclei classes.

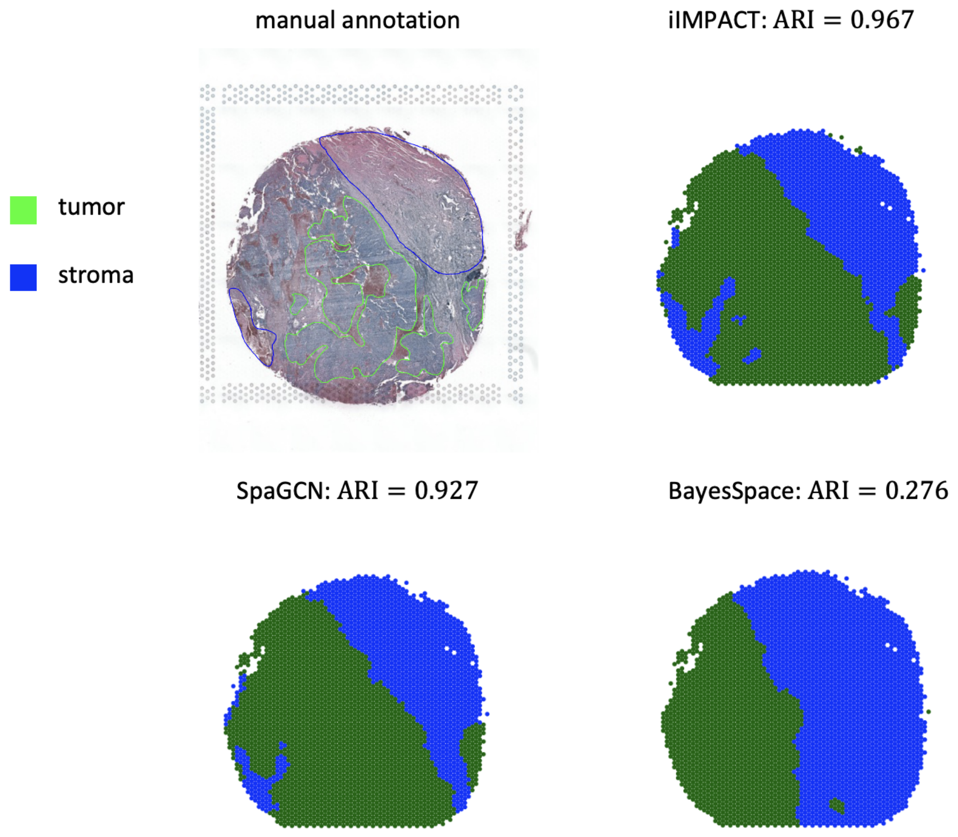


**Figure S5**: Histology image of the tissue section with manually annotated labels from pathologists, and histology-based spatial domains detected by iIMPACT, SpaGCN, and BayesSpace, setting the number of clusters to be 2, for human ovarian cancer data.

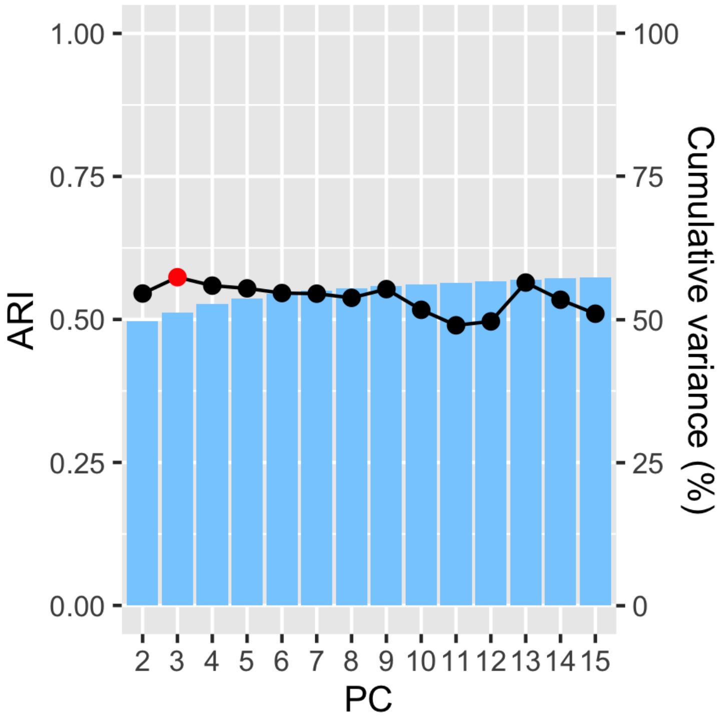


**Figure S6**: Sensitivity analysis: ARIs achieved by iIMPACT when setting the image profile weight $w$ to be zero, and proportion of variance explained under different number of leading principal components in PCA.

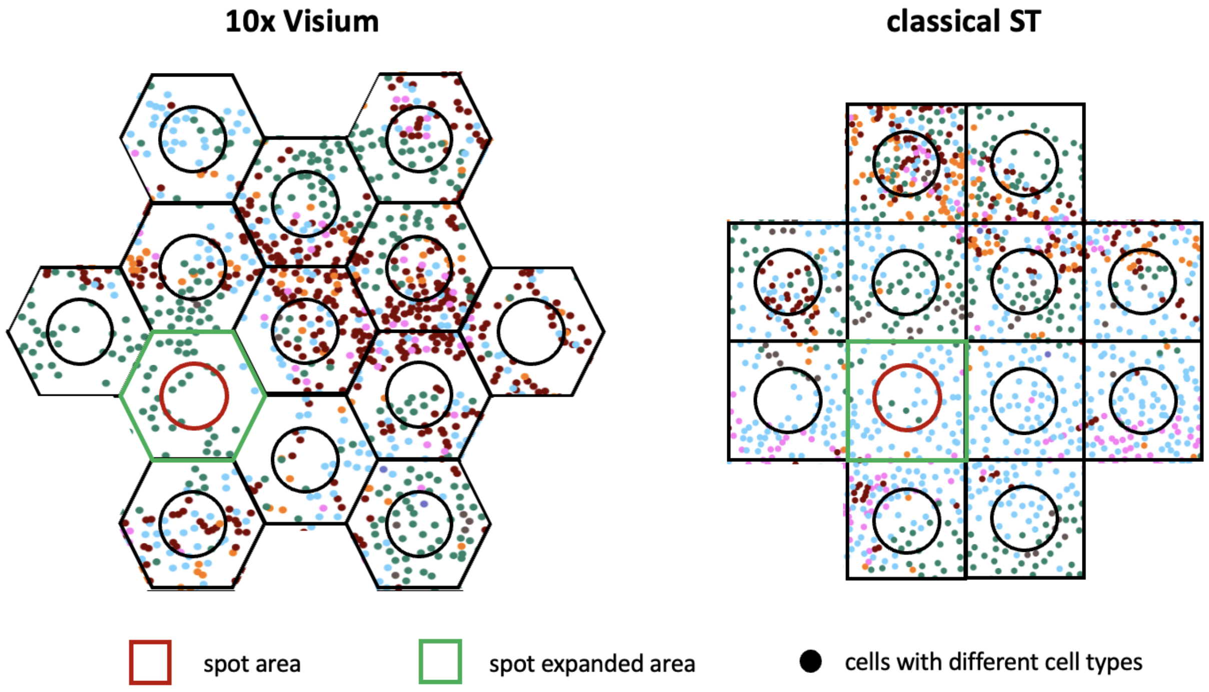


**Figure S7**: Demonstration of geometric representations of spatial distribution of spots, and definition of spot expanded area in the 10x Visium and classical ST technologies.

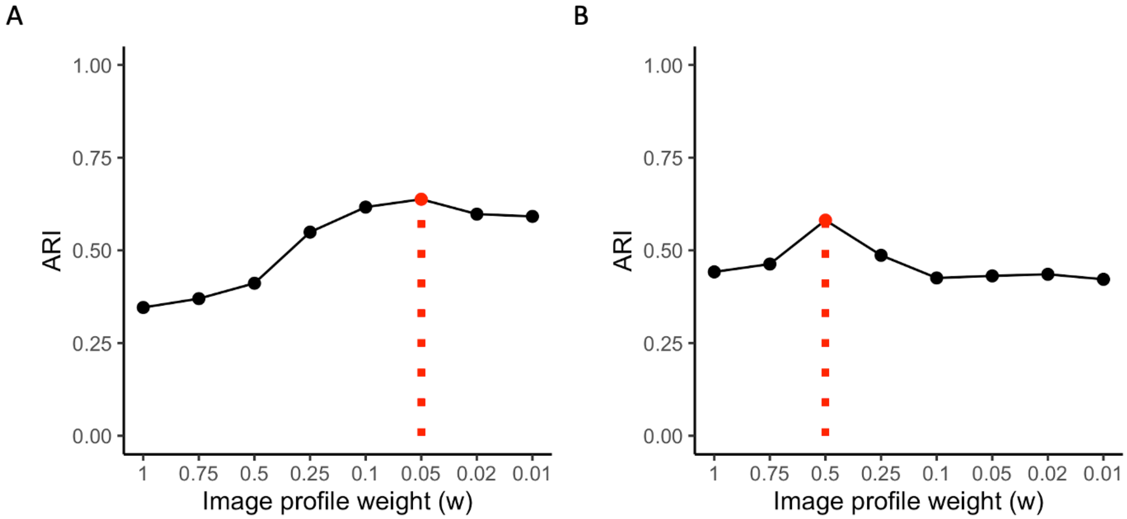


**Figure S8**: Sensitivity analysis: ARIs achieved by iIMPACT clustering under different choices of the image profile weight $w$ on A. human breast cancer data; and B. mouse visual cortex STARmap data.

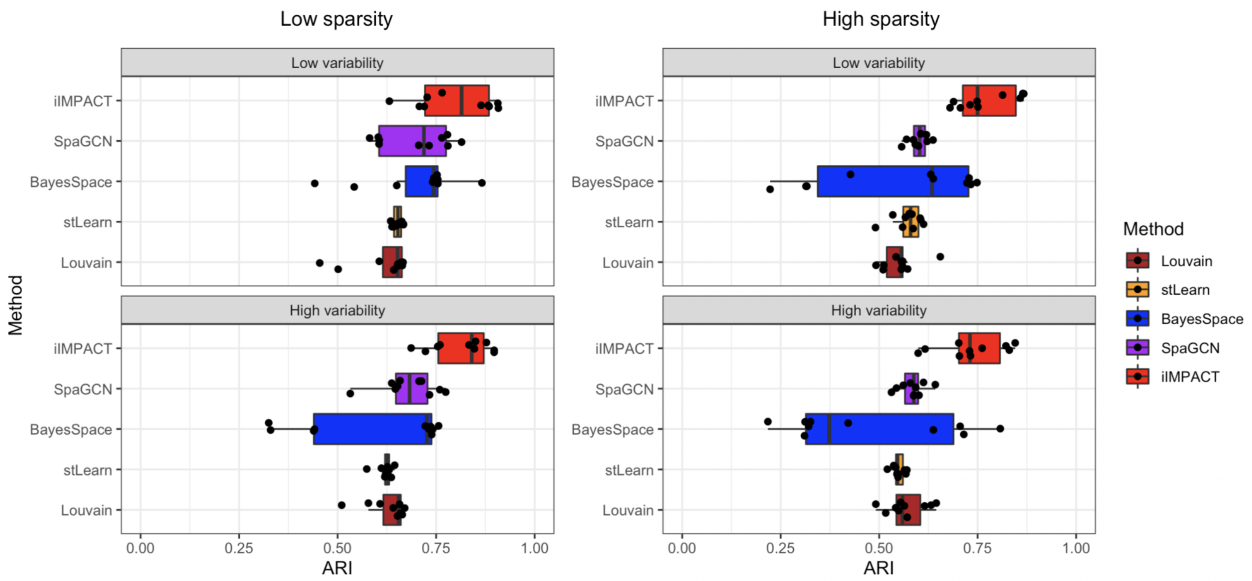


**Figure S9**: Simulation study: The boxplots of ARIs achieved by iIMPACT, SpaGCN, BayesSpace, stLearn, and Louvain under different scenarios in terms of sparsity and variability settings.

Table S1: Summary of the four real datasets analyzed in the paper.

| Dataset name | Technology | Organism | Tissue | Disease | Number of genes | Sample size | Source |
|---|---|---|---|---|---|---|---|
| Human breast cancer data | ST (10x Visium) | Human | Breast | Ductal carcinoma in situ, invasive carcinoma | 17943 | 2518 | 10x Genomics (https://www.10xgenomics.com/resources/datasets) |
| Human prostate cancer data | ST (10x Visium) | Human | Prostate | Adenocarcinoma, invasive carcinoma | 17943 | 4371 | 10x Genomics (https://www.10xgenomics.com/ |
| Human ovarian cancer data | ST (10x Visium) | Human | Ovarian | Serous papillary carcinoma | 17943 | 3455 | 10x Genomics (https://www.10xgenomics.com/resources/datasets) |
| Mouse visual cortex STARmap data | STARmap | Mouse | Brain-visual cortex | - | 1207 | 1020 | Wang et al. (2018) |

Table S2: Running time (in minutes) of iIMPACT, BayesSpace and SpaGCN on the four real datasets analyzed in the paper.

| Datasets | iIMPACT | BayesSpace | SpaGCN |
|---|---|---|---|
| Human breast cancer data | 1.78 | 18.30 | 1.83 |
| Human prostate cancer data | 2.67 | 27.99 | 2.27 |
| Human ovarian cancer data | 2.24 | 23.49 | 1.90 |
| Mouse visual cortex STARmap data | 1.09 | 13.86 | 0.57 |

Table S3: The key notations of the proposed iIMPACT.

| | Notation | Support | Definition |
|---|---|---|---|
| Data | $N$ | $N \in \mathbb{N}$ | Number of spots |
| | $P$ | $P \in \mathbb{N}$ | Number of genes |
| | $P'$ | $P' \in \mathbb{N}$, $P' \ll P$ | Number of reduced dimensions |
| | $Q$ | $Q \in \mathbb{N}$ | Number of cell types |
| | $K$ | $K \in \mathbb{N}$ | Number of histology-based spatial domains |
| | $\boldsymbol{C} = [c_{ij}]_{N \times P}$ | $c_{ij} \in \mathbb{N}$ | The gene expression count table with $c_{ij}$ being the read count for gene $j$ observed at spot $i$ |
| | $\boldsymbol{Y} = [y_{ij}]_{N \times P'}$ | $y_{ij} \in \mathbb{R}$ | The low-dimensional gene expression table with $y_{ij}$ being the relative experssion on dimension $j$ at spot $i$ |
| | $\boldsymbol{V} = [v_{iq}]_{N \times Q}$ | $v_{iq} \in \mathbb{N}$ | The cell type abundance table with $v_{iq}$ being the number of cells with cell type $q$ at spot $i$ and its expanded area |
| | $\boldsymbol{m} = [m_i]_{N \times 1}$ | $m_i \in \mathbb{N}$ | The total number of cells at spot $i$ and its expanded area |
| | $\boldsymbol{T} = [t_{ir}]_{N \times 2}$ | $t_{ir} \in \mathbb{N}$ | The $x$ and $y$ coordinates of spots |
| | $\boldsymbol{G} = [g_{ii'}]_{N \times N}$ | $g_{ii'} \in \{0, 1\}$ | The adjacent matrix with $g_{ii'} = 1$ indicating spot $i$ and spot $i'$ are neighbors |
| | $\boldsymbol{s} = [s_i]_{N \times 1}$ | $s_i \in \mathbb{R}^+$ | Size factors |
| Model parameters | $\boldsymbol{\Omega} = [\omega_{kq}]_{K \times Q}$ | $\omega_{kq} \in [0, 1]$ | With $\omega_{kq}$ being the relative abundance of cell type $q$ in histology-based spatial domain $k$ |
| | $\boldsymbol{M} = [\mu_{kj}]_{K \times P'}$ | $\mu_{kj} \in \mathbb{R}$ | With $\boldsymbol{\mu}_k$ being the mean vector of the multivariate normal subcomponent for histology-based spatial domain $k$ |
| | $\boldsymbol{\Sigma} = [\Sigma_k]_{K \times P' \times P'}$ | $\sigma_{kjj'} \in \mathbb{R}^+$ | With $\Sigma_k$ being the covariance matrix of the multivariate normal subcomponent for histology-based spatial domain $k$ |
| | $\boldsymbol{z} = [z_i]_{N \times 1}$ | $z_i \in \{1, \ldots, K\}$ | Histology-based spatial domain indicator |
| | $\boldsymbol{\psi} = [\psi_j]_{P \times 1}$ | $\psi_j \in \mathbb{R}^+$ | The gene-specific dispersion parameter |
| Tuning parameters | $w$ | $w \in [0, 1]$ | The image profile weight |

Table S4: Posterior means of parameters (domain-specific relative abundances of cell types $\hat{\boldsymbol{\Omega}}$ and means of low-dimensional representation of gene expression $\hat{\boldsymbol{M}}$) for human breast cancer data, which are applied in generating the simulated data.

| Domain-specific relative abundances of cell types $\hat{\boldsymbol{\Omega}}$ | | | | | | | |
|---|---|---|---|---|---|---|---|
| | Stromal cell | Red blood cell | Lymphocyte | Tumor cell | Karyorrhexis | Ductal epithelium | Macrophage |
| Domain 1 | 0.4209 | 0.0905 | 0.0492 | 0.3407 | 0.0415 | 0.0562 | 0.0009 |
| Domain 2 | 0.5221 | 0.1045 | 0.3346 | 0.0174 | 0.0161 | 0.0045 | 0.0007 |
| Domain 3 | 0.2551 | 0.5202 | 0.1855 | 0.0166 | 0.0141 | 0.0037 | 0.0049 |
| Domain 4 | 0.3875 | 0.2802 | 0.0498 | 0.2207 | 0.0384 | 0.0168 | 0.0066 |
| Domain 5 | 0.5327 | 0.1888 | 0.2502 | 0.0066 | 0.0203 | 0.0011 | 0.0002 |

| Domain-specific means of low-dimensional representation of gene expression $\hat{\boldsymbol{M}}$ | | | |
|---|---|---|---|
| | PC 1 | PC 2 | PC 3 |
| Domain 1 | 26.2131 | 2.3886 | -0.0389 |
| Domain 2 | -2.6583 | -5.3326 | -1.2944 |
| Domain 3 | -6.6993 | -2.6915 | 4.0473 |
| Domain 4 | -19.0341 | 10.3570 | 1.1856 |
| Domain 5 | -18.0996 | 1.3260 | -1.8454 |
